# HUH-tag conjugation enables accessible, site-specific functionalization of DNA origami with signaling-competent proteins

**DOI:** 10.64898/2026.11.11.765849

**Authors:** Alina Ehrl, Elke Ziska, Meghan A. Morrissey, Konlin Shen, Shawn M. Douglas, Marcus J. Taylor

## Abstract

DNA origami enables precise nanometer scale control over the number, spacing, and geometry of biomolecules, making it a powerful tool for dissecting spatially organized biological processes such as receptor signaling. However, functionalizing origami with proteins remains a bottleneck restricting the technology to specialized laboratories. Here we show that viral-derived HUH endonuclease tags, small (∼10 kDa) domains that form a covalent phosphotyrosine bond with short unmodified ssDNA, provide a simple and cost-effective route to decorating DNA origami with functional proteins. Using interleukin-1β (IL-1β) as a model ligand, we generated recombinant IL-1β-HUH, achieved efficient site-specific conjugation to unmodified ssDNA within minutes, and coupled the conjugates to a DNA origami pegboard with tunable stoichiometry. IL-1β-functionalized origami triggered ligand-dependent signalling in living cells and allowed visualization of the recruitment of downstream signaling molecules, confirming that HUH-tag conjugation preserves functionality. Because this strategy requires only standard molecular biology equipment and unmodified DNA, it lowers a technical and financial barrier to DNA origami-based experimentation, opening the approach to a broader range of biological questions and laboratories outside the DNA nanotechnology field.

## Introduction

Precise nanometer-scale spatial control over molecules is a defining feature of biological systems. Understanding this organization quantitatively requires tools that can interrogate and manipulate biomolecules with matching spatial precision. DNA origami provides exactly this capability by forming custom two- or three-dimensional nanostructures through folding of a long single-stranded scaffold by annealing of short DNA oligonucleotides, referred to as staple strands^1–4^. Origami nanostructures can be functionalized with biomolecules at user-defined positions, enabling independent control over number, spacing, and geometry of biomolecules at the nanometer scale ^5^. Functionalization uses staple strands extended with unique single-stranded DNA (ssDNA) sequences, which hybridize to complementary ssDNA strands fused to biomolecules^6,7^. This ability to position biomolecules, such as proteins, with nanometer precision has made DNA origami a powerful platform to study biological systems both in vivo and in vitro^7,8^, and consequently DNA origami has provided mechanistic insight into the spatial arrangement of multienzyme assemblies^9,10^, enabled structural studies by controlling protein orientation^11–13^, and revealed the molecular dynamics of receptor signalling pathways^14^. Despite this potential, DNA origami-based approaches remain largely confined to laboratories that specialise in nanofabrication and DNA nanotechnology. One technical barrier is attaching ssDNA to proteins, which relies on bioconjugation chemistry that lies outside the expertise and resources of most biology laboratories. As a result, producing functionalised DNA origami remains costly, labour-intensive and technically demanding, which limits its use in cell biology and biochemistry.

While the materials used to construct the origami (the ssDNA scaffold and staple strands) are available from commercial vendors, the essential step of linking a biomolecule to the nanostructure remains challenging. To minimize dissociation, a protein of interest should ideally be covalently attached to an ssDNA strand^7^. This typically requires the recombinant production of a protein genetically modified to contain a specific attachment site for ssDNA. Once sufficient quantities of a suitable recombinant protein are obtained, conjugation to a ssDNA label often involves complex multi-step protocols. Existing protein-DNA conjugation strategies each involve trade-offs. Site specific labelling methods that obtain a 1:1 stoichiometry fuse a protein of interest to genetically encoded self-labeling enzyme tags, such as SNAP-tag^15^ and HaloTag^16^, or peptide tags that can be labelled enzymatically. Self-labelling enzymes are relatively large (20 kDa or larger), potentially compromising the function of the protein. In contrast, peptide tags that are recognized by enzymes such as Sortase A^17^ are smaller but require optimization of slow and reversible labeling conditions to achieve sufficient yields of ssDNA-protein conjugate^18,19^. Non-specific chemical conjugation methods, such as NHS-ester or maleimide coupling to native amino acids, are faster and cheaper bypassing the need to fuse a tag to the protein of interest, but sacrifice site-specificity and one-to-one stoichiometry, and yield heterogeneous, non-quantitative conjugation. Non-specific conjugation can also compromise the bioactivity of a protein^20,21^. Finally, the modified oligonucleotides needed for all of these conjugation methods must either be purchased at considerable cost or made in laboratories equipped for DNA synthesis and organic chemistry. No current ssDNA-protein conjugation method therefore combines site-specific, stoichiometric labelling with low cost and ease of use, and this has limited the broader adoption of DNA origami within the cell biology and biochemistry community.

Here, we present a straightforward, cost-effective strategy for decorating DNA origami with recombinant proteins using HUH endonuclease tags, so called because of the presence of two conserved histidines (H) separated by a hydrophobic residue (U)^22^. Viral-derived HUH tags are small (approx. 11 kDa) domains that form a covalent phosphotyrosine linkage to a short, unmodified ssDNA recognition sequence (15 nt for the tag used here), thus enabling site-specific, high-yield conjugation under physiological buffer conditions. This removes the need for purchasing or generating complex chemically modified oligonucleotides, and allows protein-DNA conjugates to be generated without expert knowledge in organic chemistry synthesis, with significant reductions in time and cost. To demonstrate the applicability of this approach, we apply this strategy to functionalize a previously described DNA origami pegboard^23,24^ with interleukin-1β (IL-1β), a signaling system that to our knowledge has not been previously investigated with DNA origami. We generate recombinant HUH-tagged IL-1β, establish the labeling conditions with ssDNA attachment sequences, and demonstrate precise control over stoichiometry on the origami surface. Finally, we demonstrate that this conjugation strategy preserves protein function by using IL-1β-functionalized origami to trigger IL-1 receptor signaling in living cells. This establishes HUH-tag conjugation as a broadly applicable strategy for functionalizing DNA origami with active biomolecules, implemented using the standard equipment found in biochemical and cell biological laboratory settings.

## Results

### HUH-tags as an efficient tool for production of ssDNA-labelled IL-1β

An ideal strategy for labeling proteins with ssDNA is a covalent attachment chemistry that is rapid (within an hour reaction time), can proceed in a single reaction step, can be performed at physiological conditions to maintain bioactivity, and minimizes costly specialized ssDNA reagents that must be produced ahead of time or purchased from specialized vendors. We reasoned that the best way to achieve this would be a small (less than 20 kDa), highly soluble tag that could be easily genetically fused to a protein of interest for recombinant production. In principle, HUH endonucleases derived from viruses fulfil these requirements and have been shown to covalently attach single-stranded DNA to proteins ^22^. To test whether HUH endonucleases could functionalize DNA origami, we fused a previously characterized HUH endonuclease to the C-terminus of recombinant IL-1β^25^. We selected IL-1β as it has been found to tolerate C-terminal fusions^26^ yet had not been previously conjugated to DNA or DNA origami. Therefore, this fusion to IL-1β provided a suitable test case for implementation of this strategy.

Size exclusion chromatography of purified IL-1β-HUH revealed a single predominant peak consistent with a soluble, monomeric fusion protein (Fig. S2). We next assessed whether the HUH tag retained its DNA-conjugation activity when fused to IL-1β. Recombinant IL-1β-HUH was incubated with 35-nucleotide ssDNA oligonucleotides containing the 15-nucleotide HUH recognition sequence, and conjugation was monitored by SDS-PAGE over 1 h (Fig. 1A, B). A shift in apparent molecular weight consistent with formation of the protein-DNA conjugate indicated successful coupling. Optimization of the reaction conditions yielded conjugation efficiencies of approximately 70% within 10 min, as estimated by densitometric analysis of Coomassie-stained gels (Fig. 1B). These results demonstrate that the HUH tag retained conjugation activity following fusion to IL-1β, enabling efficient covalent DNA attachment. This is consistent with HUH tag offering an economical solution for generating protein-DNA conjugates for cell biological and biochemical applications.

**Figure 1.**
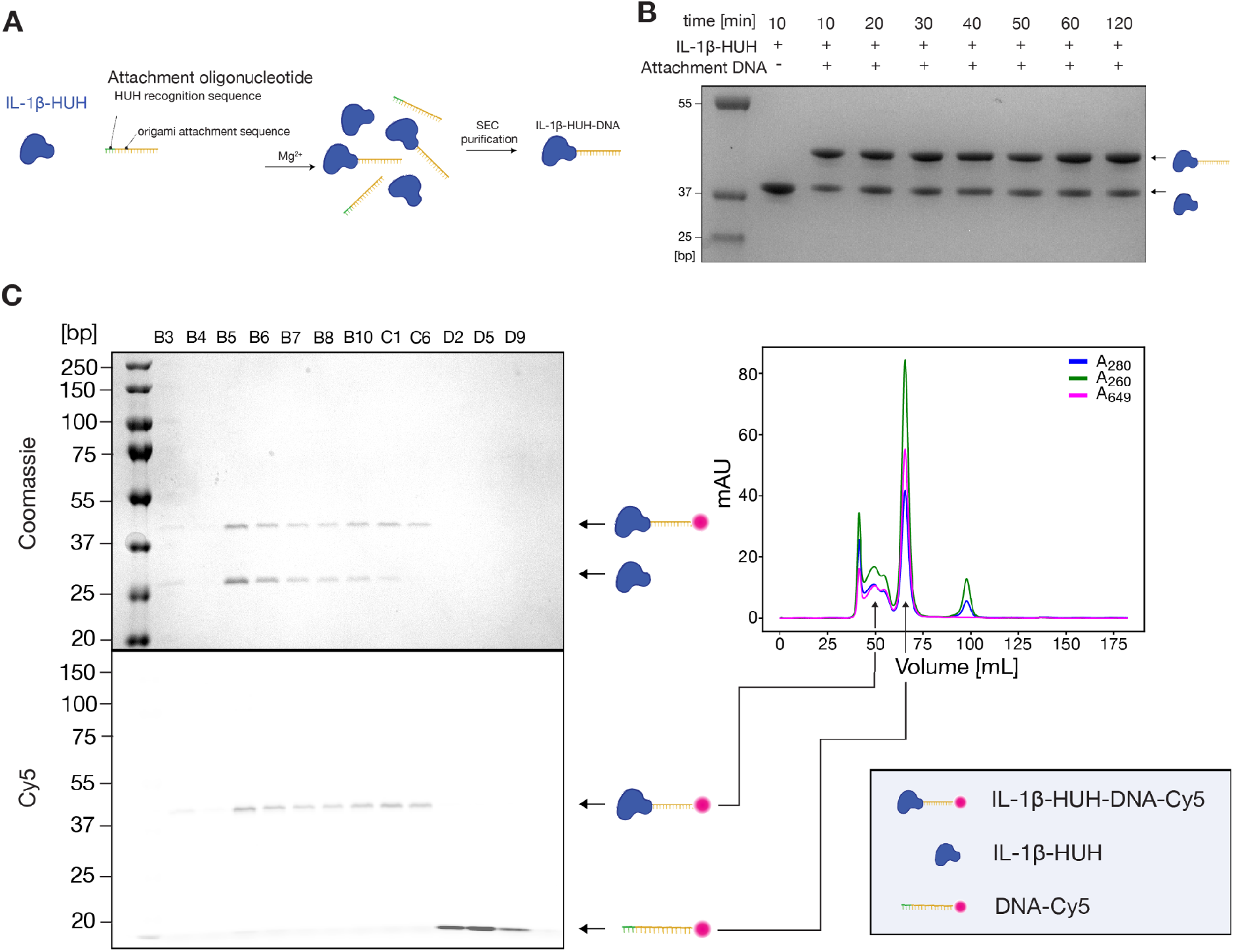
Site-specific conjugation of IL-1β to DNA using a HUH endonuclease tag. **(A)** Strategy for site-specific protein-DNA conjugation. IL-1β-HUH reacts with single-stranded DNA containing the HUH recognition sequence, forming a covalent protein-DNA conjugate. The resulting IL-1β-HUH-DNA conjugates can subsequently be purified by size-exclusion chromatography (SEC) to remove unreacted components. **(B)** Time course of labelling of IL-1β with single-stranded attachment DNA. Representative Coomassie-stained SDS-PAGE gel. Bands corresponding to IL-1β-HUH and the IL-1β-HUH-DNA conjugates are indicated. **(C)** Separation of free Cy5-conjugated DNA from protein-DNA-Cy5 conjugates. SDS-PAGE analysis of the purification of the DNA-labelled ligand. Coomassie-stained gel of fractions of size exclusion chromatography run (top left), fluorescence scan of SDS-PAGE gel of respective fractions (bottom left) and SEC chromatogram (right).

Investigating receptor signaling using DNA origami requires controlled protein loading to minimize variation in ligand number between nanostructures. Protein-DNA conjugates bind to complementary attachment strands on the origami, but residual unconjugated oligonucleotides can compete for these sites and reduce loading efficiency. We therefore established a size-exclusion chromatography protocol to separate IL-1β-HUH-DNA conjugates from unreacted oligonucleotides. Purification was assessed using fluorophore-labelled DNA and fluorescence detection following SDS-PAGE, which showed separation of free oligonucleotides from the purified conjugate fraction (Fig. 1C). This demonstrates that HUH tags could be used to generate a homogenous preparation of protein-DNA conjugate for downstream application with standard protein purification equipment routinely found in biochemical and cell biology labs. Importantly bioconjugation and purification of the conjugate can be achieved within one day. Together, these results demonstrate a cost-efficient and specific strategy for generating purified IL-1β-DNA conjugates suitable for controlled attachment to DNA origami scaffolds.

### Site-specific decoration of origami nanostructures with ssDNA-conjugated ligands

Having established robust yields of IL-1-DNA conjugates, we next asked whether these conjugates could be used to generate a functionalized DNA origami platform. To maximize accessibility and avoid the overhead of de novo design, we selected a previously validated DNA origami “pegboard” (Fig. S1A through C) that had been used to study Fc receptor and T cell receptor signaling^23,24^, and thus offered design features well suited to probing the spatial organization of cell surface receptors. This design had not, however, previously been functionalized with a protein ligand, providing a stringent test of whether it could support spatially controlled presentation of HUH-tagged proteins. Origami folding was performed using a commercially available ssDNA scaffold together with ssDNA staple strands modified to be complementary to the DNA sequences conjugated to IL-1-HUH. Folding reactions were carried out on a thermocycler, purified by PEG precipitation and the resulting structures were assessed by negative stain transmission electron microscopy, confirming correct assembly into the expected three-dimensional geometry (Fig. S1D and E). These results establish that an off-the-shelf DNA origami design can be folded correctly and functionalized with attachment sequences compatible with HUH-tagged proteins, a key step toward broadening the accessibility of this platform to the cell biology and biochemistry community.

Having confirmed correct folding of DNA origami bearing ssDNA attachment sequences complementary to those fused to IL-1b-HUH, we next sought to generate IL-1-functionalized origami. Precise, nanoscale control over ligand presentation requires two conditions: high occupancy of the ssDNA tethers by conjugated ligand, and near-complete removal of unbound protein, together yielding a homogeneous population of fully loaded origami. We addressed both requirements with a thermal annealing protocol^6^ in which IL-1β-HUH-DNA conjugates were hybridized to complementary tethers on the origami surface via a controlled temperature ramp, followed by purification using size-exclusion chromatography (Fig. 2A). Chromatographic analysis with a Superose 6 column resolved two clearly separated peaks corresponding to DNA origami and free protein-DNA conjugate, with no detectable overlap, indicating efficient removal of unbound IL-1β-HUH-DNA from the reaction mixture. To confirm that protein-DNA conjugates were physically associated with the origami structures, IL-1β-HUH was labeled with a Cy5-conjugated oligonucleotide prior to hybridization; fluorescence detection during SEC showed that the Cy5 signal co-eluted precisely with the origami peak, confirming successful hybridization of ligand to scaffold (Fig. 2B). Together, these results demonstrate that a previously established DNA origami structure can be functionalized with an HUH-tagged protein of interest, providing a cost-effective and experimentally streamlined route for deploying this technology in a cell biological context.

**Figure 2.**
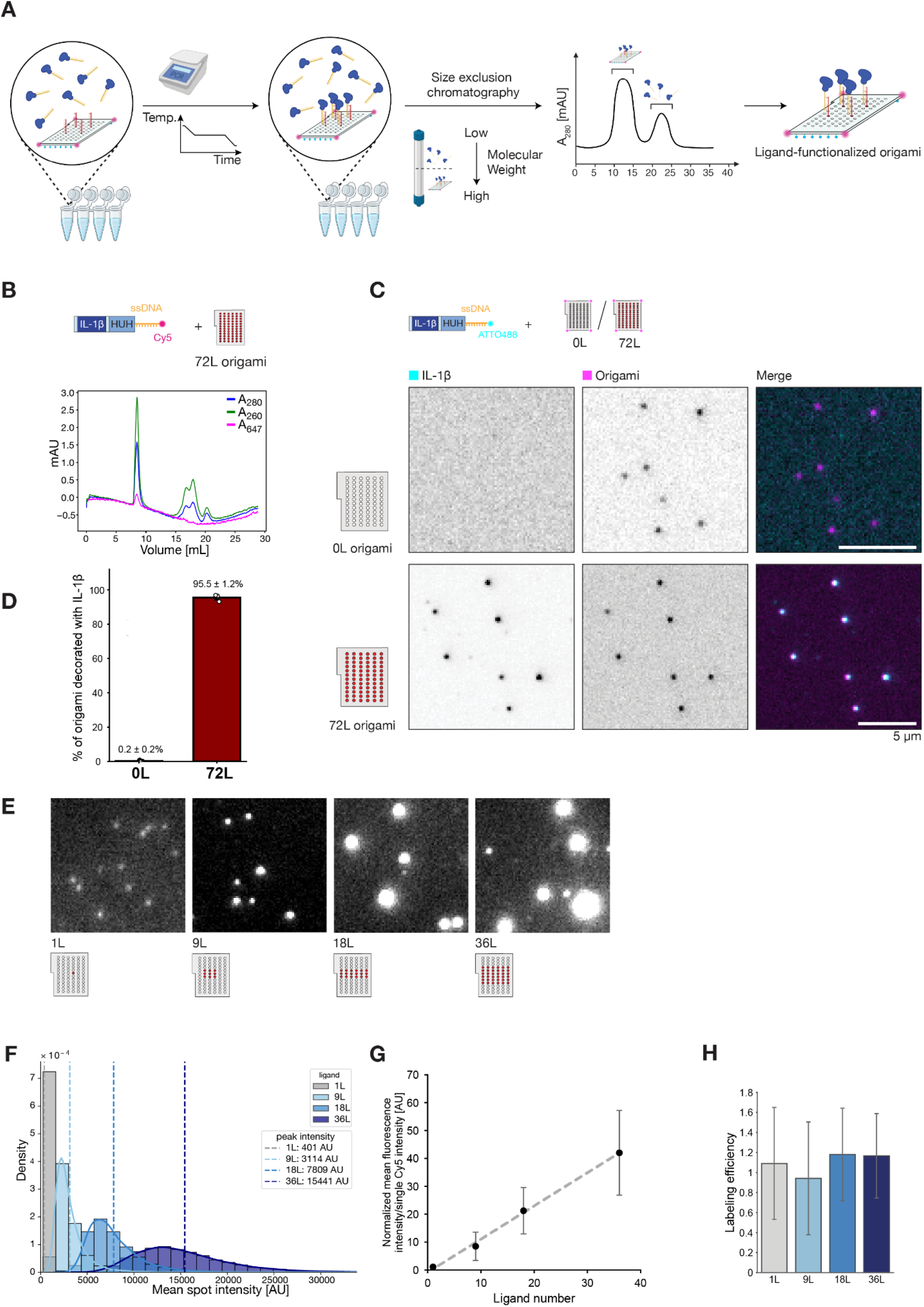
Specific and quantitative functionalization of DNA origami scaffolds with IL-1β via site-directed hybridization. **(A)** Schematic of the functionalization and purification workflow. IL-1β-HUH-DNA conjugates were hybridized to complementary attachment tethers on DNA origami via a controlled thermal annealing ramp, then separated from unbound conjugates by size-exclusion chromatography (SEC) to yield homogeneously loaded, ligand-functionalized origami. **(B)** SEC chromatograms of origami presenting 72 IL-1β-HUH-DNA-Cy5 ligands (72L), monitored by absorbance at 260 nm, 280 nm and 647 nm. Two resolved peaks correspond to DNA origami (early fraction) and free protein-DNA conjugate (late fraction). **(C)** Representative two-color TIRF microscopy images of ATTO488-labelled IL-1β-HUH-DNA and ATTO647N-labelled DNA origami for tether-free (0L) and 72 ligand (72L) origami designs, with merged overlays (IL-1β-HUH-DNA-ATTO488: cyan, Origami-ATTO647N: magenta). Scale bar: 5 μm. **(D)** Fraction of DNA origami-ATTO647N puncta associated with IL-1β-HUH-DNA-ATTO488 for 0L (grey) and 72L (red) origami designs. Data are mean ± s.d., n = 3 replicates (points). **(E)** Representative TIRF images of Cy5-labelled IL-1β-HUH loaded onto origami carrying 1, 9, 18 or 36 attachment sites (1L-36L). **(F)** Fluorescence-intensity distributions of individual DNA origami bearing 1, 9, 18 or 36 ligand attachment sites. Solid curves show Gaussian fits; vertical dashed lines indicate the corresponding peak intensities. **(G)** Estimated ligand occupancy as a function of designed ligand number, determined by normalizing origami fluorescence intensities to the intensity of a single Cy5 fluorophore. The dashed line indicates a linear fit. Data are mean ± s.d. (n = 3). **(H)** Ligand-loading efficiency, calculated as the ratio of estimated ligand occupancy to designed ligand number, with 1 corresponding to complete occupancy. Data are mean ± s.d. (n = 3).

### Single-particle analysis confirms stoichiometric labelling and homogenous preparation of functionalized DNA origami

Many applications of DNA origami in cell signaling have combined this technology with high-resolution microscopy, enabling nanoscale molecular control to be leveraged for the study of dynamic biological systems. Because such applications rely on the detection, visualization and analysis of single origami structures, often interacting with live cells, they require highly homogeneous preparations of functionalized origami to support robust biological conclusions, for instance regarding the role of receptor clustering in signaling^14^. We therefore turned to microscopy to assess the homogeneity of our IL-1b-functionalized DNA origami.

To first establish the specificity of ligand attachment, we performed two-color TIRF microscopy using ATTO647N-labeled DNA origami and IL-1β-HUH conjugated to ATTO488-labeled DNA, comparing a control structure lacking exposed attachment tethers (0L) with a design containing 72 attachment tethers on the origami surface (72L). In the absence of attachment tethers, no ATTO488 signal was detected at the positions of origami puncta (Fig. 2C), indicating that the ligand does not bind non-specifically to the origami surface. By contrast, the attachment-competent design showed strong colocalization between the ATTO488 ligand signal and the ATTO647N origami signal (Fig. 2C), and quantification confirmed a substantial increase in colocalized puncta relative to the control (Fig. 2D). Together, these results demonstrate that IL-1β-HUH-DNA binding is strictly dependent on complementary attachment sequences, establishing the site-specificity of our hybridization strategy and confirming a key prerequisite for generating homogeneous samples suitable for cell biological applications.

Having established the specificity of the hybridization reaction, we tested whether this strategy could be extended to exploit the stoichiometric and spatial control afforded by DNA origami, by generating a series of homogeneous origami bearing defined numbers of protein-DNA conjugates. A panel of origami designs containing increasing numbers of ligand attachment sites (1-36 ligands) was generated and hybridized with Cy5-labeled IL-1β-HUH-DNA conjugates to enable fluorescence-based quantification (Fig. 2E). Individual origami particles were imaged by TIRF microscopy to assess sample homogeneity, revealing diffraction-limited puncta with uniform size and intensity distributions and minimal aggregation, consistent with well-defined, monomeric origami structures. To estimate ligand occupancy, the mean fluorescence intensity of individual puncta was measured for each design; histograms of spot intensities across all conditions revealed distinct, well-separated distributions (Fig. 2F), consistent with homogeneous origami populations bearing discrete numbers of IL-1-HUH-DNA conjugates. Stoichiometry was estimated by fitting Gaussian functions to each intensity distribution and dividing the resulting mean intensity by the fluorescence intensity of single Cy5 fluorophores, revealing close agreement between expected and measured labeling stoichiometry across the entire range (Fig. 2G). Converting these values to labeling efficiency showed that all origami designs, regardless of target labeling number, achieved a near-complete labeling efficiency (Fig. 2H).

Together, these results demonstrate that the developed hybridization and purification workflow enables efficient and specific functionalization of DNA origami scaffolds with IL-1β, while allowing quantitative control over ligand number at the single-particle level, a prerequisite for systematic studies of biological systems^7,27^. We conclude that HUH tags can be combined with DNA origami to create a flexible, cost-effective experimental platform, offering the wider biological community a means to exploit the nanoscale spatial and stoichiometric control afforded by DNA origami.

### IL-1β-functionalized origami triggers intracellular molecular dynamics characteristic of productive IL-1 receptor signaling

An important consideration in developing DNA origami to interrogate biological systems is that functionalized proteins retain their biological activity once conjugated to the origami structure. Having established efficient and specific functionalization of DNA origami with IL-1β-HUH, we next sought to determine whether IL-1β-HUH-functionalized origami retained functional activity. One key advantage of DNA origami for studying receptor signalling is that fluorescent origami presenting a defined number of ligands can be combined with live-cell fluorescence microscopy, allowing the recruitment of fluorescently tagged signalling proteins to be visualised at sites of receptor engagement^14,23,24^, offering unique insight into how signaling pathways operate at the molecular level. Productive IL-1 signaling is defined by the formation of the multimeric Myddosome signaling complex at the IL-1 receptor and subsequent recruitment of downstream signaling proteins to this complex^26,28^. Therefore, we tested whether our functionalized DNA origami could trigger these molecular dynamics, thereby providing a high-resolution validation that our origami platform could stimulate the molecular dynamics of functional IL-1 receptor signaling in live cells, and that HUH-tag conjugation did not perturb IL-1b activity.

To visualize the molecular dynamics by TIRF microscopy, DNA origami boards containing 72 possible ligand attachment sites were hybridized with Cy5-conjugated IL-1β-HUH-DNA and immobilized on glass surfaces (Fig. 3A). This configuration enabled spatially defined presentation of IL-1β ligands within the evanescent field of the TIRF microscope. To monitor signaling events downstream of IL-1 receptor activation, we used EL4 cells, in which IRAK1 was endogenously tagged at its C-terminus with the bright green fluorescent protein StayGold, and NEMO was endogenously tagged at its N-terminus with mScarlet-i (Fig. 3A, B and S2, S3). EL4 cells were selected as a model system because they overexpress IL-1 receptors and respond robustly to IL-1β stimulation. Within the IL-1β signaling cascade, IRAK1 represents the final kinase recruited during Myddosome assembly and thus serves as a marker of fully assembled Myddosomes, whereas NEMO acts downstream of the Myddosome as part of the IKK complex, linking Myddosome formation to signal propagation toward NF-κB activation. Visualizing both of these molecular events at IL-1b-functionalized origami would therefore confirm that IL-1β-HUH retains functional activity.

**Figure 3.**
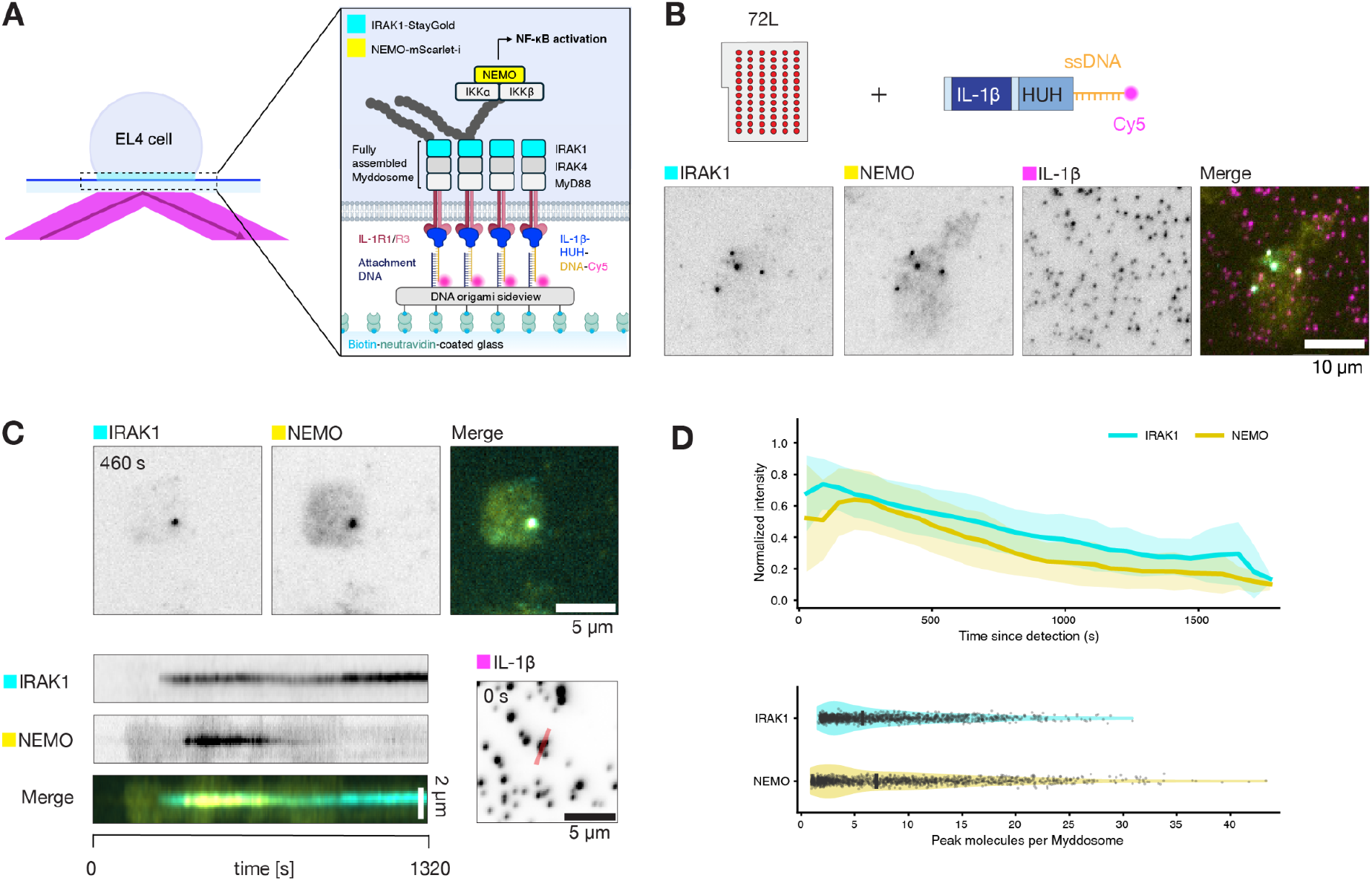
DNA origami-presented IL-1β induces Myddosome assembly and downstream NEMO recruitment. **(A)** Schematic of the TIRF microscopy assay. EL4 cells expressing IRAK1-StayGold and mScarlet-i-NEMO interact with IL-1β–functionalized DNA origami immobilized on biotin-neutravidin-coated glass. Fluorescently tagged IRAK1 and NEMO report Myddosome assembly and recruitment of the downstream IκB kinase (IKK) complex, respectively. When exposed to EL4 cells, IL-1 receptors bind to the presented ligands, enabling visualization of signaling complex formation within the TIRF illumination field. **(B)** Representative TIRF microscopy images of EL4 cells interacting with IL-1β-functionalized DNA origami boards after 30 min of stimulation. Endogenously tagged IRAK1-StayGold (cyan) and mScarlet-i-NEMO (yellow) accumulate in puncta that colocalize with Cy5-labeled IL-1β-decorated origami particles (magenta). Representative fields of view and merged channels are shown. Scale bar: 10 μm. **(C)** Time-resolved recruitment of signaling components to ligand-functionalized origami. Representative frames and kymographs showing the accumulation of IRAK1-StayGold (cyan) and mScarlet-i-NEMO (yellow) at an individual IL-1β-decorated 72L origami particle (magenta). The red line indicates the position used for the kymograph analysis. Scale bar: 5 μm (top), 2 μm (bottom). **(D)** Recruitment dynamics and molecular abundance of IRAK1 and NEMO. Top, mean fluorescence-intensity trajectories aligned to initial detection and normalized to the maximum intensity of each track in the corresponding channel. Solid lines and shaded regions indicate mean ± s.d. Bottom, distributions of peak IRAK1 and NEMO molecule numbers per Myddosome, estimated using single-fluorophore intensity calibrations. Violin plots show the distributions; individual points represent tracked assemblies.

Upon contact between cells and ligand-functionalized origami surfaces, both IRAK1 and NEMO accumulated into discrete puncta that colocalized with Cy5-labeled origami particles (Fig. 3B, C), demonstrating that functional signaling complexes assemble specifically at sites of ligand presentation. Control experiments confirmed that EL4 cells did not respond to origami structures lacking IL-1β, indicating that the observed signaling events are ligand-dependent rather than a consequence of nonspecific interactions with the DNA nanostructure (Fig. S3). Time-resolved imaging further revealed that signaling dynamics on the origami platform recapitulate the established temporal hierarchy of IL-1 receptor signaling: IRAK1 was recruited first and formed stable puncta (Fig. 3C), followed by subsequent recruitment of NEMO, consistent with the known sequential assembly of the Myddosome and downstream activation of the IKK complex. Together, these results demonstrate that IL-1β presented on DNA origami scaffolds is capable of triggering Myddosome formation and localized receptor signaling, with the recruitment dynamics of IRAK1 and NEMO indicating that the origami-based platform faithfully reproduces key molecular features of IL-1 signaling while enabling spatially controlled ligand presentation. We conclude that HUH tags provide a cost-effective and accessible strategy for generating functionalized DNA origami to interrogate dynamic biological systems such as signaling pathways.

To extend our observations beyond individual events, we quantified the recruitment dynamics of IRAK1 and NEMO across the full population of detected puncta. Averaged intensity traces revealed that IRAK1 and NEMO followed a consistent temporal pattern across many independent recruitment events, with IRAK1 intensity rising rapidly upon detection and NEMO recruitment following in close succession, before both signals gradually declined over the course of the imaging window (Fig. 3D). This population-level analysis confirmed that the sequential recruitment pattern observed for individual puncta (Fig. 3C) reflects a general feature of signaling complex assembly at IL-1β-functionalized origami rather than an isolated event. To estimate the stoichiometry of signaling complexes formed at the origami surface, we quantified the peak number of IRAK1 and NEMO molecules recruited per punctum. Both proteins displayed broad distributions in peak molecule number, consistent with heterogeneity in Myddosome and IKK complex assembly at the single-particle level, with NEMO spanning a wider range than IRAK1 (Fig. 3D). Together, these analyses demonstrate that IL-1β-functionalized origami reproducibly trigger the ordered recruitment of IRAK1 and NEMO across hundreds of individual ligand-presenting particles. By combining stoichiometrically defined, site-specific protein conjugation with single-particle imaging in live cells, we demonstrate that HUH-tags are an effective strategy to conjugate ssDNA to a protein of interest which can be used to create functionalized DNA origami to examine cell biological processes such as receptor activation.

## Discussion

Here, we established HUH endonuclease tags as a simple and cost-effective way to functionalize DNA origami nanostructures with recombinant proteins. Using a C-terminal HUH-fusion tag, we achieved covalent, site-specific conjugation of IL-1β to short, unmodified ssDNA oligonucleotides at high labelling efficiencies under physiological buffer conditions, without the need for chemically modified oligonucleotides or specialist conjugation chemistry. We further showed that these protein-DNA conjugates could be hybridized onto DNA origami pegboards with high specificity and tunable stoichiometry, ranging from 1 to 36 ligands per particle, and that this functionalized origami preserved IL-1β bioactivity, as demonstrated by ligand-dependent sequential recruitment of IRAK1 and NEMO at the site of origami engagement. Together, these results establish HUH-tag conjugation as a practical route to construct protein-functionalized DNA origami nanostructures using only equipment and reagents already standard in most molecular biology and biochemistry labs.

A central motivation for this work was accessibility. Conventional strategies for protein-DNA conjugation, such as self-labelling enzyme tags (e.g. SNAP-tag and HaloTag) or sortase-mediated ligation, each require a large fusion domain (32 kDa in the case of HaloTag), slow and low-yielding reaction chemistry, or chemically modified oligonucleotides carrying reactive handles (e.g. benzylguanine, chloroalkane or oligoglycine). These modified oligonucleotides are expensive and only available from a limited number of specialist suppliers. By contrast, HUH tags react with unmodified ssDNA that can be ordered from any oligonucleotide vendor. For the 35-nt attachment oligonucleotide used here, the unmodified sequence costs approximately $12 per nmol, approximately six-fold less than the equivalent benzylguanine-modified oligonucleotide purchased ready to use (approximately $74 per nmol; Supplementary Table 9). To save cost, the benzylguanine substrate can be prepared in-house by coupling a 5’-amine-modified oligonucleotide to BG-GLA-NHS, but this adds chemical conjugation and purification steps. The BG-GLA-NHS reagent alone represents an upfront outlay of over $500. Either route therefore imposes a cost or an additional chemistry step that HUH conjugation removes entirely. Combined with a conjugation and purification workflow that can be completed within a single day using standard IMAC, SEC and SDS-PAGE equipment, this substantially lowers the barrier for non-specialist laboratories to generate functionalised DNA origami.

The HUH endonuclease superfamily is functionally and structurally diverse, and different members recognize distinct ssDNA sequences^22^. Among these, the viral rolling-circle replication initiator proteins are particularly well suited as fusion tags: they are small, and they target short unmodified ssDNA sequences (15 nt for the FBNYV-derived tag used here). In contrast, the bacterial relaxase-type HUH endonucleases, which have previously been used to assemble proteins orthogonally on DNA origami nanostructures^29^, are considerably larger and recognize substantially longer ssDNA targets^25^. Within the viral subfamily, orthogonal tag/oligonucleotide pairs have already been identified, and deep sequencing of target-sequence libraries has resolved sufficiently differences in specificity to extract orthogonal sets, with two- and three-way orthogonal labelling validated experimentally and supported by a dedicated substrate design tool^25,30,31^. Self-labelling tags such as SNAP-tag and HaloTag, by contrast, offer only a small number of mutually orthogonal chemistries^15,16^. The small size of the viral HUH tags (approx. 11 kDa, compared with approx. 20 and 32 kDa for SNAP-tag and HaloTag respectively) further reduces the risk that the tag perturbs folding, solubility or function of the protein of interest, consistent with our observation that IL-1β-HUH remained monomeric, soluble and functionally active as a receptor ligand. Together, these properties suggest that HUH tags could be extended to generate multi-component DNA origami platforms in which several distinct proteins are independently and orthogonally conjugated to defined positions on the same nanostructure, for example a ligand and a competing antagonist, or the reconstitution of multiple components of signaling or metabolic complexes. The complexity of such platforms would then be set by the origami design rather than by the available conjugation chemistry.

Finally, the ability of IL-1β-functionalized origami to trigger the expected temporal hierarchy of IL-1 receptor signal transduction (IRAK1 recruitment followed by NEMO) in live cells confirms that HUH conjugation does not compromise receptor-ligand functionality and that DNA origami can be used to interrogate the spatiotemporal dynamics of a signaling pathway at the single complex level (Fig. 3C, D). We therefore anticipate that this strategy, combining HUH-tag based protein-DNA conjugation with off-the-shelf origami designs, will be readily transferable to other receptor-ligand pairs, and more broadly to any application requiring precise stoichiometric and site-specific presentation of proteins on DNA nanostructures, including in vitro reconstitution of multi-enzyme assemblies and mechanistic studies of substrate channeling. HUH-tag conjugation reduces the functionalization of DNA origami to a single-step protein biochemistry reaction, placing the nanoscale control afforded by these nanostructures within reach of any laboratory able to purify a recombinant protein.

## Methods

### Cell culture

Wild-type EL4.NOB1 cells (ECACC; hereafter referred to as EL4) and their gene-edited derivatives were maintained at 37 °C under 5% CO₂ in RPMI medium (Thermo Fisher Scientific) containing 10% FBS (Biozol) and 2 mM L-glutamine. Cultures were passaged every 2–3 days to maintain cell densities of 0.1-0.5 × 10⁶ cells/mL. Mycoplasma testing with the MycoAlert detection kit confirmed that all cell lines were negative.

### Generation of CRISPR/Cas9 engineered cell lines

EL4 cells were transfected with sgRNA/Cas9-expressing pX330 plasmids and pMK-based HDR templates using the Neon Transfection System. Electroporation was performed with a single 50-ms pulse at 1,080 V. For simultaneous editing of two loci, sgRNA/Cas9 and HDR template plasmids were combined at equal molar ratios, using 1.5 μg of plasmid DNA. Cells were subsequently cultured in antibiotic-free RPMI medium for 24 h before puromycin was added at 1μg/mL. Selection was maintained for 48 h.

Monoclonal lines were established by fluorescence-activated cell sorting (FACS) using a BD FACSAria II at the Flow Cytometry Core Facility of the Deutsches Rheuma-Forschungszentrum Berlin. An initial bulk sort enriched cells positive for both fluorescent tags. Following expansion, individual cells were sorted into 96-well plates containing culture medium supplemented with 15% EL4-conditioned RPMI medium.

Clonal lines were characterized by PCR, Sanger sequencing, and western blotting. Genomic DNA was extracted using QuickExtract DNA Extraction Solution (Epicentre). Integration of the mStayGold and mScarlet-i cassettes was assessed by PCR across the junctions linking the fluorescent protein coding sequences, the homology arms, and the target loci. Additional PCR assays spanning the tag cassette, an entire homology arm, and the adjacent genomic sequence were used to distinguish heterozygous from homozygous clones. Amplification products were resolved on 0.8-1% agarose gels, purified using Monarch Nucleic Acid Purification Kits (NEB), and submitted for Sanger sequencing. These analyses indicated homozygous editing at both the IRAK1 locus and the NEMO locus in the EL4 IRAK1-mStayGold/mScarlet-i-NEMO cell line.

Expression of the tagged proteins was assessed by Western blotting with antibodies against IRAK1 and NEMO (see Table 1 for antibody details). Fluorescent tagging increased the apparent molecular weight by approximately 25 kDa relative to the corresponding untagged proteins. All clones were additionally examined by fluorescence microscopy to assess the subcellular localization of the tagged proteins.

### Homology-directed repair (HDR) DNA template design for CRISPR/Cas9 endogenous labeling of NEMO and IRAK1

Homology-directed repair (HDR) templates for tagging IRAK1 with monomeric StayGold (mStayGold)^32^ and NEMO with mScarlet-i^33^ were constructed in a pMK vector backbone (Life Technologies, Carlsbad, CA), following a strategy previously described^26^. Homology arms amplified from EL4 genomic DNA were combined with fragments encoding the respective fluorescent tag and in the case of NEMO an upstream puromycin resistance cassette, separated by a 2A sequence^34^. Silent mutations were introduced into the homology arms to disrupt sgRNA target sites, protecting the repair templates from Cas9 cleavage. All fragments were incorporated into the pMK backbone by Gibson Assembly, and the resulting plasmids were verified by sequencing. Details of individual HDR template constructs are provided below.

#### pMK-PuroR-2A-mScarlet-i-NEMO HDR

The NEMO HDR template contained 785-bp and 890-bp homology arms flanking the ATG start codon of the mouse NEMO gene (ENSMUSG00000004221). A PuroR–2A-mScarlet-i cassette was positioned between the arms with an in frame 3x(Gly-Gly-Ser) linker connecting NEMO and the fluorescent tag. The following primer pairs were used for amplification: 5′ homology arm: forward 5′-TTGTAAAACGACGGCCAGCTCTTTTATCATGTGTTTGTGTTTTCTTGACTCC-3′ and reverse 5′-CATCCAACAGGACAGTGTCTAGAAGAAAAGGGTAG-3′; 3′ homology arm: forward 5′-GGAAACAGCTATGACCATGAAGAAGTCAAACCAATTCCTAGCCAGTTC-3′ and reverse 5′-TGGAAGAACCAGCTGAGTGAGATGG-3′. PuroR–2A–mScarlet-i–cassette: forward 5′-AGACACTGTCCTGTTGGATGACCGAGTACAAGCCCACGG-3′ and reverse 5′-CACTCAGCTGGTTCTTCCAAGGGTGCTTGTTACTGCCTCCTGAGCCACCCTTGTA-3′; pMK vector backbone: forward 5′-CTGGCCGTCGTTTTACAA-3′ and reverse 5′-CATGGTCATAGCTGTTTCC-3′. PCR fragments were combined and fused using Gibson assembly. The full-length constructs were confirmed by Sanger sequencing.

#### pMK-IRAK1-mStayGold-HDR

To construct a HDR template to insert a mStayGold ORF at the C-terminus of IRAK1, we modified a previously described HDR plasmid^26^. This template contained 2251-bp and 764-bp homology arms flanking the stop codon of the mouse IRAK1 gene (ENSMUSG00000031392). The following primer pair was used to amplify the HDR arms and pMK vector backbone: forward 5’-TCTGAGACACTTGAGGCCCACCTCTGATTTGTTCACTCTGACAAATCCCT-3’ and reverse 5’-AAACAGTTCTTCTCCTGTACTAACACTACCACCAGAACCTCCACTTCCTCC-3’. A mStayGold ORF codon optimized cassette was ordered as a gBlock from IDT that contained a 3x(Gly-Gly-Ser) linker connecting IRAK1 and mStaygold open reading frame. The gBlock and PCR product were fused using Gibson Assembly and the full-length construct was confirmed by Sanger sequencing.

### Generation of CRISPR/Cas9 sgRNA vectors

Single-guide RNAs (sgRNAs) targeting sites within 50 bp upstream or downstream of the NEMO N-terminal coding sequence were designed using the Benchling CRISPR design tool. Complementary oligonucleotides for each guide were purchased from Integrated DNA Technologies (IDT) and cloned into pX330 (pX330-U6-Chimeric_BB-CBh-hSpCas9; Addgene #42230), which expresses *Streptococcus pyogenes* Cas9 and a chimeric guide RNA. The oligonucleotides were 5′-phosphorylated with T4 polynucleotide kinase, annealed, and ligated into BbsI-digested pX330 using Quick Ligase (NEB). The resulting plasmids were transformed into NEB Stable competent cells, and all sgRNA constructs were verified by sequencing. Of the NEMO guides tested, one sgRNA generated the most positive cells by FACS and was selected to generate cell lines expressing tagged NEMO. For endogenous labelling of IRAK1, a previously described sgRNA was used^26^. Details of sgRNA sequences are given in Table 1.

### Cloning of the pET28a-His10-IL-1β-HUH expression construct

To functionalize individual origami nanostructures with IL-1β ligands via ssDNA tethers on the origami, we created a recombinant mature active form of murine IL-1β_(118-269)_ which was C-terminally fused to a viral HUH endonuclease from FBNYV allowing for specific protein-ssDNA conjugation. The sequence encoding for recombinant His_10_-IL-1β-HUH was codon optimized for expression in *E. coli* using the OPT webserver^35^ and ordered as gBlocks from IDT. The HUH-tag was fused to IL-1β via a 3xGGGS linker. Using Gibson assembly^36^, the gene fragment was cloned into the SpeI-HF/NheI-HF digested pET-28a-IL-1β-spytag plasmid (previously described here^26^). The seamless joining of adjacent fragments is achieved through complementary overlapping ends of ∼20 bp which were included into the design of the gBlock. The digested empty pET-28a backbone was purified through gel extraction using the NEB Monarch DNA Gel Extraction Kit and for the Gibson Assembly, 50 ng of digested vector were combined with the gBlock insert at a 1:3 molar ratio to the vector. The DNA mixture was combined with NEBuilder HiFi DNA Assembly Master Mix at a 1:3 volume ratio, incubated at 50 °C for 1 h and transformed into chemically competent NEB 10-beta *E. coli* cells following the manufacturer’s instructions. The cells were plated on agar plates containing 100 μg/mL ampicillin and incubated at 37 °C overnight. Colonies were randomly selected and screened by colony PCR using the DreamTaq Hot Start Green PCR Master Mix and a T7 promotor forward primer and a T7 terminator reverse primer. Positively screened colonies were inoculated in LB media containing 100 μg/mL ampicillin and plasmid DNA was extracted using the NEB Monarch Plasmid Miniprep Kit according to the manufacturer’s instructions (pET-28a-His_10_-IL-1β-HUH).

### Cloning of the pET28a-His_10_-mScarlet-i expression construct

The sequence encoding for N-terminally His_10_-tagged mScarlet-i^33^ was codon-optimized for expression in *E. coli* and ordered as gBlocks from IDT. The sequences were cloned into the pET-28a backbone via SpeI-HF/NheI-HF enzyme digestion and Gibson assembly by M. J. Taylor (pET-28a-His_10_-mScarlet-i).

### Cloning of the pET28a-His_10_-mStayGold expression construct

The sequence encoding for N-terminally His_10_-tagged monomeric StayGold^32^ was codon-optimized for expression in *E. coli* using the Twist Codon Optimization Tool and cloned into the pET-28a vector by Twist Bioscience (pET-28a-His_10_-StayGold).

### Protein purification

All proteins were expressed in chemically competent BL21 (DE3) *E. coli* cells. Protein expression was induced at OD_600_ = 0.8 - 1.2 using 250 µM IPTG and cultures were grown overnight at 18 °C. The purification process for each protein was monitored using SDS-PAGE.

For purification of His_10_-IL-1β-HUH as well as His_10_-StayGold, cells were lysed in lysis buffer 1 (50 mM HEPES pH 7.5, 200 mM NaCl, 5 mM imidazole, 2 mM MgCl_2_, 1X protease inhibitor, 1X DNaseI, 1X lysozyme, 1 mM DTT) using ultrasonication, respectively. After clearing the lysate, both constructs were purified via immobilised metal affinity chromatography using Ni-NTA resin in HisTrap wash buffer 1 (50 mM HEPES pH 7.5, 250 mM NaCl, 20 mM imidazole, 1 mM DTT) and eluted in HisTrap elution buffer 1 (50 mM HEPES pH 7.5, 250 mM NaCl, 500 mM imidazole, 1 mM DTT). Peak fractions were pooled and concentrated, followed by gel filtration using a Superdex 200 10/300 column equilibrated in SEC buffer 1 (20 mM HEPES pH 7.5, 150 mM NaCl). Respective peak fractions were concentrated, snap-frozen and stored at -80 °C.

His_10_-mScarlet-i was purified following the purification steps described above using lysis buffer 2 (50 mM HEPES pH 8, 200 mM NaCl, 5 mM imidazole, 2 mM MgCl_2_, 1X protease inhibitor, 1X DNaseI, 1X lysozyme, 1 mM DTT), HisTrap wash buffer 2 (50 mM HEPES pH 8, 250 mM NaCl, 20 mM imidazole, 1 mM DTT), HisTrap elution buffer 2 (50 mM HEPES pH 8, 250 mM NaCl, 500 mM imidazole, 1 mM DTT) and SEC buffer 2 (20 mM HEPES pH 8, 150 mM NaCl), respectively.

### SDS-PAGE

To evaluate protein composition, fresh or previously frozen samples that were stored at -20 °C were used at room temperature and mixed with a final concentration of 1X loading dye and subsequently boiled for 5 min at 95 °C. SDS-PAGE was performed using either NuPAGE 4-12% Bis-Tris protein gels or discontinuous gels with a 10% separating gel in Bolt Empty Mini Gel Cassettes (Laemmli 1970). Samples were loaded onto the gels and run in either 1X MES running buffer or 1X Laemmli buffer (Laemmli 1970) at 160 V for 15 min, then at 180 V for 40 min. Proteins were visualized using InstantBlue Ultrafast Protein Stain.

### Protein-DNA conjugation

For conjugation of an attachment ssDNA sequence to HUH-tagged proteins, different oligonucleotides were ordered from IDT at HPLC-grade purity, all containing the *ori* recognition sequence of the FBNYV-derived HUH tag at the 5’ end: 5’-AAGTATTACCAGAAA-3’. The recognition sequence was followed by an attachment sequence complementary to variable attachment handles on the origami. For all experiments on IL-1β-HUH, we used the attachment sequence A: 5’-CCTAGGGTATTGAATGAGGG-3’. To introduce additional fluorescent labeling of either the ligand or the origami, fluorophore-conjugated versions of the labeling oligonucleotides were ordered from IDT at HPLC-grade purity by adding an additional 3xT linker to the 3’ end, followed by either Cy5 or ATTO488.

To conjugate His_10_-IL-1β-HUH with ssDNA, the protein was mixed at 5 to 10 μM with a 2X excess of the labeling oligonucleotide in 0.5X HUH buffer (25 mM HEPES pH 8, 25 mM NaCl, 0.5 mM MgCl2) with additional 1 mM MgCl_2._ The mixture was incubated at 37 °C for 30 min.

The labeling reactions were applied to a HiLoad 16/600 Superdex 75 gel filtration column equilibrated in SEC buffer 1 (20 mM HEPES pH 7.5, 150 mM NaCl), the peak fractions were pooled and concentrated. The labeling efficiency was estimated by SDS-PAGE. To evaluate the removal of excess oligonucleotide, the labeling reaction was performed using a Cy5-conjugated attachment oligonucleotide and subjected to purification via size exclusion chromatography as described above. The peak fractions were analyzed by SDS-PAGE and fluorescently labelled oligonucleotides were visualized using the LI-COR Odyssey CLx Infrared Imaging System. The protein-ssDNA conjugates were snap-frozen and stored at -80 °C.

### DNA origami

We used a DNA origami design that was previously developed to study T cell activation and FcγR-mediated phagocytosis^23,24^. The attachment sequences were modified to achieve higher DNA-DNA affinities for stable functionalization of the origami structures with a protein ligand as well as to provide the possibility of hybridization of multiple ligand species. The 8064-nt-long single-stranded scaffold DNA (type p8064) was ordered from tilibit from a M13mp18 derivative at a concentration of 100 nM. DNA staple oligonucleotides were ordered from IDT, purified by standard desalting and resuspended in water at 100 μM in 96-well plates. ATTO647N-conjugated and biotinylated oligonucleotides were ordered to HPLC-grade purity from IDT. The list of all oligonucleotides used for the different origami designs in this study are listed in Table 2-5.

The origami nanostructures were self-assembled in a mixture containing 20 nM scaffold DNA and 200 nM staple DNA in origami folding buffer FoB_20_ (5 mM Tris pH 8, 5 mM NaCl, 1 mM EDTA, 20 mM MgCl_2_), incubated at 65 °C for 15 min, then annealed with a temperature ramp from 60 °C to 20 °C with a decrease in temperature of 1 °C/h.

To remove excess staple DNA, the origami particles were purified by PEG precipitation by carefully mixing an equal volume of PEG precipitation buffer (15% (w/v) PEG-8000, 5 mM Tris-Base pH 8.0, 1 mM EDTA, 500 mM NaCl, 20 mM MgCl_2_) with an equal volume of DNA origami folding reaction followed by centrifugation at 16000 × g for 25 min at room temperature. The supernatant was removed and the translucent pellet resuspended in folding buffer FoB_15_ (5 mM Tris pH 8, 5 mM NaCl, 1 mM EDTA, 15 mM MgCl_2_). The folded origami structures were stored at 4°C protected from light.

### Agarose gel electrophoresis

The success of the assembly of the origami was evaluated by agarose gel electrophoresis using 2% agarose gels in 1X TAE (40 mM Tris base, 40 mM acetate, 1 mM EDTA), 0.5 μg/mL Gel Red stain and 11 mM MgCl_2_. The gels were run at 80 V for approximately 2 h in an ice-cold water bath and then imaged using the GelDoc XR+ gel documentation system.

### Negative stain transmission electron microscopy

Before sample application, 400-mesh hex grids (or higher mesh size) coated with a thin carbon film were glow discharged for 60 s. 5 μL of approximately 100 ng/μL origami sample in origami folding buffer FoB_15_ (5 mM Tris pH 8, 5 mM NaCl, 1 mM EDTA, 15 mM MgCl_2_) were applied to the grids with an absorption time of 1 min before blotting. The grid was washed by dipping into a 15 μL droplet of origami folding buffer FoB_15_ and subsequent blotting. This washing step was repeated before dipping the grid into a 15 μL droplet of 2% aqueous uranyl formate or 2% aqueous uranyl acetate, blotting followed by dipping the grid into another 15 μL stain droplet for 45 s and subsequent blotting. Before imaging, the grids were air-dried for a minimum of 15 min to overnight.

### Hybridization of DNA-conjugated protein to DNA origami

To achieve a site-specific decoration of the DNA origami nanostructures with functionalized proteins, 1 pmol of origami with *n* attachment handles was combined with a 5X excess per *n* attachment handles of DNA-conjugated protein in 20 mM HEPES pH 7.5 and 15 mM MgCl_2_. The sample was incubated at 37 °C for 1h, then cooled to 22°C with a decrease of 0.1 °C/min, followed by incubation at 22 °C for 14 h and a decrease in temperature of 0.1°C/min until a temperature of 4 °C was reached. To remove excess protein, the sample was gel-filtrated using a Superose 6 Increase 10/300 GL column equilibrated in origami SEC buffer (20 mM HEPES pH 7.5, 150 mM NaCl, 15 mM MgCl_2_). The peak fractions corresponding to protein-decorated origami nanostructures were pooled and directly used for further experiments.

### Coupling of origami nanostructures to glass

For cleaning of the imaging plate, each well in a 96-well glass bottom plate was filled with cleaning solution containing 5% Hellmanex III and 10% isopropanol and incubated at 50 °C for 30 min on a heating plate. The plate was emptied, filled with fresh 5% Hellmanex III and incubated at 50 °C for 1 h on a heating plate. Subsequently, the plate was thoroughly washed with dH_2_O at least 20 times, dried off with argon gas and sealed with a plate sealer or directly used for experiments. To passivate the glass surface, each well was incubated overnight at 4 °C with 0.5 mg/mL Biotin-BSA in HBS, freshly prepared from a 10X stock solution and enough volume to cover the entire bottom of the well, followed by incubation at room temperature for 1 h. Each well was washed with 10 mL 1X HBS before adding 0.05 mg/mL neutravidin, enough to cover the entire bottom of the well and freshly prepared from a 10X stock by dilution in HBS. After 2 h incubation, each well was washed with 10 mL of 1X HBS with 15 mM MgCl_2_. Serial dilutions of either the non-functionalized or the protein-decorated origami structures were created to find concentrations where individual origami spots can be resolved by TIRF microscopy. The diluted origami samples were added to the passivated and biotin/neutravidin-coated wells to fill the bottom of each well and incubated at room temperature for 15 min. Each well was washed with 10 mL of 1X HBS with 15 mM MgCl_2_. Unless indicated otherwise, all TIRF experiments were prepared as described above.

### TIRF microscopy data acquisition

Imaging of mScarlet-i-NEMO and IRAK1-StayGold recruitment was performed on an inverted microscope (Nikon TiE) equipped with a Nikon fiber-coupled TIRF illuminator. Excitation was provided by a laser combiner using 488 nm, 561 nm and 640 nm laser lines at approximately 0.35 mW, 0.25 mW, and 0.17 mW, respectively, measured at the objective output.

Fluorescence emission was collected using bandpass filters for GFP (525 ± 25 nm), RFP (595 ± 25 nm), and Cy5 (700 ± 75 nm). Images were acquired with a Nikon Plan Apo 100×/1.4 NA oil-immersion objective and projected through a 1.5× intermediate magnification lens onto a Photometrics Prime 95B sCMOS camera. Images were recorded with 2 × 2 pixel binning, resulting in a final pixel size of 150 nm.

Image acquisition was controlled using NIS-Elements software. All experiments were conducted at 37 °C using an OKO Labs stage-top incubation enclosure to maintain temperature. Images were acquired at intervals of 1-4 s with exposure times ranging from 60 to 300 ms.

### TIRF live cell imaging

Origami-functionalized wells were prepared as described above. Prior to cell addition, wells were imaged by TIRF microscopy to verify a uniform distribution of individual origami spots and comparable surface densities across experimental conditions.

To enable quantitative fluorescence analysis and single-molecule intensity calibration, calibration images were acquired before each imaging experiment using recombinant monomeric His_10_-StayGold and His_10_-mScarlet-i proteins, as well as ATTO647, Cy5, and ATTO488 dyes. For single-molecule calibration, purified monomeric His_10_-StayGold was diluted in HBS and adsorbed onto KOH-cleaned glass using 96-well glass bottom plates. Individual fluorophores were imaged using identical microscope settings to those applied during live-cell acquisition. The same procedure was performed for His_10_-mScarlet-i and the respective dyes. All calibration samples were diluted to concentrations that ensured spatially separated diffraction-limited single-molecule detection by TIRF microscopy.

For live-cell imaging, EL4 cells were added to wells coated with BSA-biotin and neutravidin and functionalized with IL-1β-decorated DNA origami. EL4 cells expressing IRAK1-StayGold and mScarlet-i-NEMO were sequentially illuminated with a 488 nm laser (60 ms exposure, 10% power), a 561 nm laser (300 ms exposure, 2% power) and a 640 nm laser (200 ms exposure, 30% power) at a frame interval of 4 s.

For cell-free experiments assessing IL-1β binding specificity to origami tethers or quantifying origami loading states, images were acquired at frame intervals of 1-4 s with exposure times ranging from 60-300 ms.

### Image analysis

Microscopy image analysis was done in Fiji 1.54p. Diffraction-limited and time-resolved puncta corresponding to IRAK1-StayGold, mScarlet-i-NEMO, origami-ATTO647, IL-1β-HUH-DNA-Cy5 or IL-1β-HUH-DNA-ATTO488 were detected and tracked using the TrackMate 8.1.6 plugin in Fiji^37^. The Fiji kymograph plugin was used for kymograph visualizations.

To quantify ligand loading on DNA origami structures, fluorescence intensity calibration was performed at the single-molecule level. Single fluorophores were imaged under identical acquisition settings as experimental samples, and diffraction-limited spots were detected and quantified using TrackMate. The resulting intensity distributions were exported and analyzed in Python. A Gaussian model was fitted to the distribution of single-fluorophore intensities to determine the mean fluorescence intensity μ_single_, representing the characteristic signal of an individual fluorophore. In parallel, fluorescence intensities of ligand-functionalized DNA origami were quantified using the same detection and analysis pipeline (μ_origami_). The distribution of origami-associated spot intensities was similarly fitted with a Gaussian function to extract the mean intensity. The average loading state of individual origami structures was then estimated by normalizing μ_origami_ to μ_single_, yielding the effective number of fluorophores per origami. Loading efficiency was calculated by comparing the measured ligand number to the theoretical maximum number of available attachment sites per origami.

Individual cells within each field of view were outlined manually on the time-averaged projection of the mScarlet-i-NEMO channel using the Fiji ROI Manager. Each tracked Myddosome was assigned to the cell whose region of interest (ROI) contained its (x,y) centroid at first detection; puncta not contained within any drawn ROI were excluded from per-cell analyses. Segmentation and tracking results were verified for every field of view by visual inspection of a quality-control overlay image showing cell outlines and the initial position of every tracked Myddosome superimposed on a maximum-intensity projection of the IRAK1 channel.

For population-level kinetic analysis, the intensity trace of each Myddosome in the IRAK1 and NEMO channels was independently normalized to that track’s own peak (maximum) intensity in the corresponding channel, yielding normalized traces ranging from 0 to 1. Normalized traces were aligned to the frame of first detection, and relative time was divided into 30 equal-width bins spanning the observed track durations; normalized intensities were averaged across all tracks within each bin to generate population-averaged recruitment curves.

The peak (maximum) spot intensity reached by each Myddosome in the IRAK1 and NEMO channels was used as a proxy for the maximal amount of each protein recruited. To convert peak intensities into estimated molecule numbers, the mean fluorescence intensity of a single StayGold and a single mScarlet molecule was determined as described above under imaging conditions identical to those used for live-cell acquisition, yielding mean single-molecule intensities of 177.8 AU (StayGold) and 139.6 AU (mScarlet). Peak spot intensities were divided by the corresponding single-fluorophore intensity to obtain the estimated number of IRAK1 and NEMO molecules recruited per Myddosome. All downstream analysis was performed in Python 3.11.0 using pandas 3.0.1 , NumPy 2.4.2 , SciPy 1.17.1 and Matplotlib 3.10.8 . Visualization of data was performed using custom scripts in Python 3.11.0.

### Statistical Analysis

All data are expressed as the mean ± standard deviation (s.d.) or the standard error (SEM), as stated in the figure legends and results. Sample sizes (number of independent experiments, fields of view, cells and tracked Myddosomes) are reported in the corresponding figure legends.

## Acknowledgements

We thank Fakun Cao (MPIIB) for assistance with TIRF microscopy. We acknowledge Volker Brinkmann and Christian Goosmann from the MPIIB Microscopy Core Facility for providing help with electron microscopy data acquisition and sample preparation. We thank Iain Patten and all Taylor lab members for critical feedback and commenting on the project.

## Funding

This work was supported by the Max Planck Society (M.T.). M.A.M. was supported by the National Institutes of Health (R35GM14693). S.D. and K.S were supported by the National Institutes of Health (R35GM125027, awarded to S.D).

## Author contributions

M.J.T. and A.E. conceived the study. A.E. designed and cloned all protein constructs, purified and labelled the proteins, and performed DNA origami folding, functionalization optimization, TIRF microscopy assays and western blotting. M.J.T. designed the CRISPR knock-in cell line, and E.Z. generated and validated it. M.A.M., K.S. and S.M.D. designed and provided the DNA origami platform. M.A.M. provided experimental protocols and K.S. provided technical guidance. A.E. analysed the data. A.E. and M.J.T. wrote the manuscript.

## Data availability

The image analysis scripts are available on GitHub (https://github.com/MJ-Taylor-Lab/Origami_Paper).

## Notes

### Competing Interest Statement

M.J.T. is a scientific advisory board member at RheumaGen.

