## Supplemental Figures for "HUH-tag conjugation enables accessible, site-specific functionalization of DNA origami with signaling-competent proteins"

### Contents:

**Figure S1 - 3**

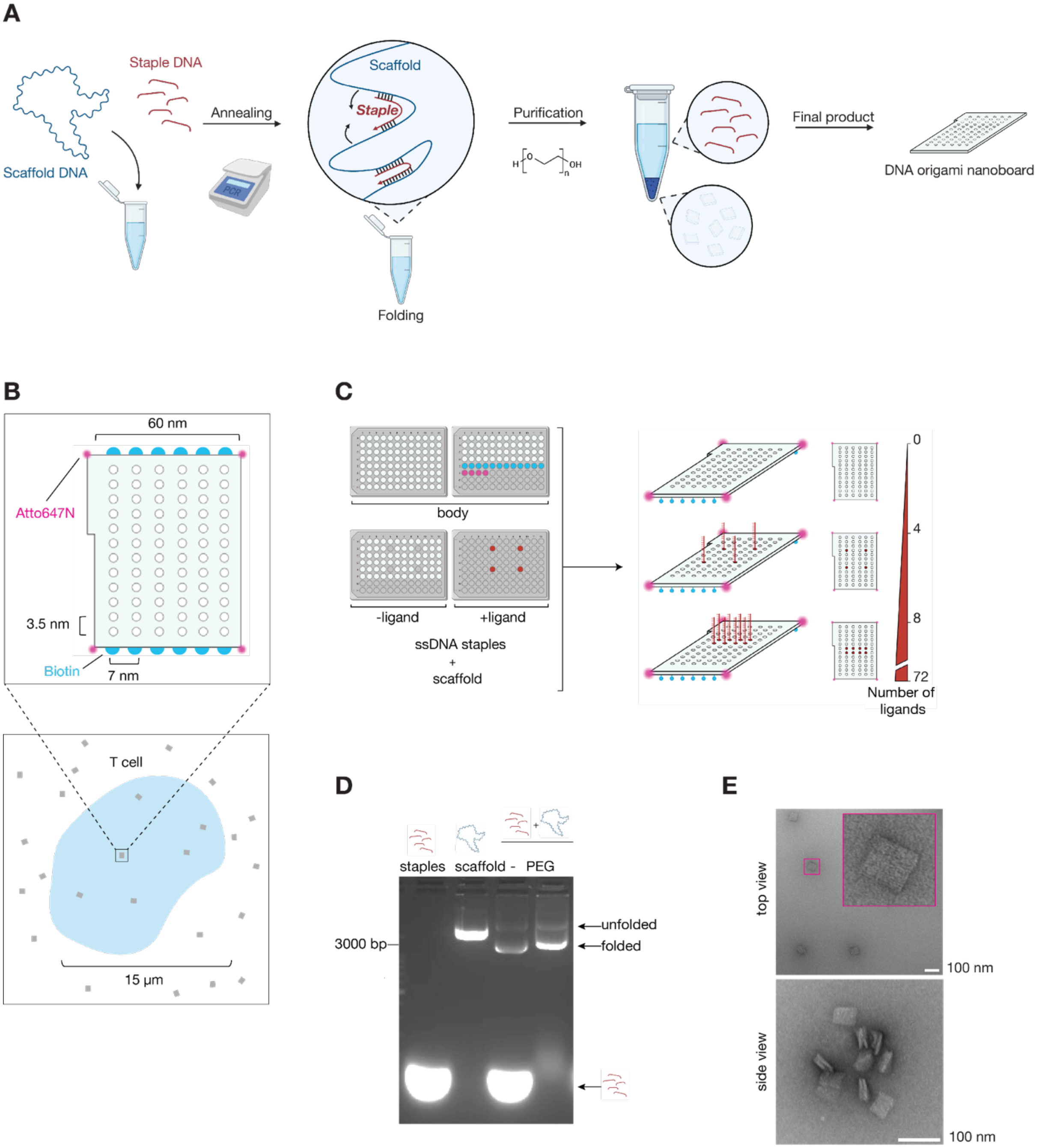


**Figure S1 | Design, assembly and characterization of a programmable DNA origami platform for ligand presentation.**

**(A)** Schematic of DNA origami assembly and purification. A single-stranded DNA scaffold is folded by complementary staple strands during thermal annealing. Polyethylene glycol (PEG) precipitation separates folded origami from excess staples.

**(B)** Schematic of the approximately 60 × 60 nm DNA origami nanoboard, comprising 72 addressable positions arranged in a 6 × 12 array with intersite spacings of 7 and 3.5 nm along the two axes. ATTO647N fluorophores (magenta) enable visualization, and biotin modifications (cyan) mediate surface immobilization. The lower schematic illustrates surface-presented origami beneath a T cell.

**(C)** Modular staple design enables control of ligand number and spatial arrangement. Selected staples are extended with single-stranded DNA handles (red) for ligand attachment, whereas unmodified staples occupy the remaining positions. Combining these staples with the scaffold and invariant structural staples generates origami with up to 72 ligand attachment sites.

**(D)** Agarose gel electrophoresis of staple strands, scaffold DNA and assembled origami before (−) and after PEG purification. The indicated bands correspond to unfolded scaffold, folded origami and excess staples.

**(E)** Negative-stain transmission electron micrographs showing top and side views of the DNA origami. The inset shows an enlarged view of the particle outlined in magenta. Scale bars, 100 nm.

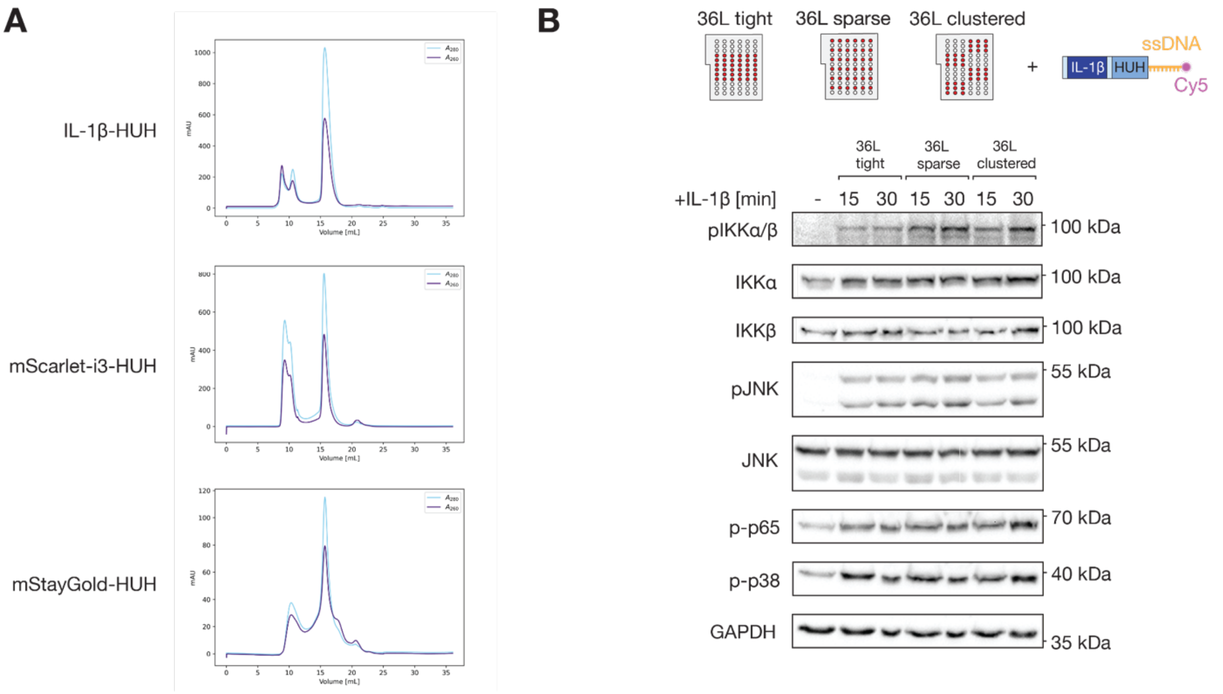


**Figure S2 | HUH-tagged proteins are soluble and origami-presented IL-1β-HUH retains signalling activity.**

**(A)** Size-exclusion chromatography profiles of IL-1β-HUH (top), mScarlet-i3-HUH (middle) and mStayGold-HUH (bottom), monitored by absorbance at 280 nm (light blue) and 260 nm (purple). Each fusion protein exhibits a major elution peak consistent with a soluble, monomeric species.

**(B)** Immunoblot analysis of EL4 cells stimulated for 15 or 30 min with DNA origami presenting 36 IL-1β-HUH ligands in tight, sparse or clustered arrangements. Unstimulated cells (−) serve as a control. Phosphorylated IKKα/β, JNK, p65 and p38 are shown alongside total IKKα, IKKβ and JNK, with GAPDH as a loading control.

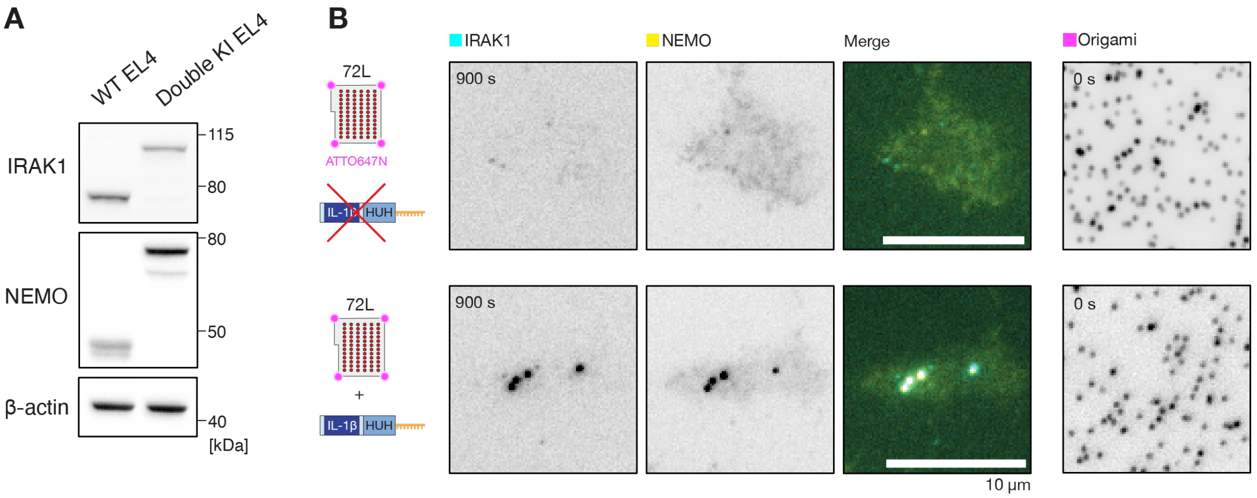


**Figure S3 | Origami-bound IL-1β induces IRAK1 and NEMO puncta formation in a dual-tagged EL4 reporter cell line.**

**(A)** Immunoblot analysis of IRAK1 and NEMO in wild-type (WT) and dual-knock-in (double KI) EL4 cells to validate fluorescent tagging of both proteins. Molecular masses are indicated in kDa.

**(B)** Representative TIRF microscopy images of EL4 cells expressing endogenously tagged IRAK1–StayGold (cyan) and NEMO–mScarlet-i (yellow) on surface-immobilized DNA origami bearing 72 ligand attachment sites, either without ligand (top) or functionalized with IL-1β-HUH (bottom). Scale bars, 10 µm.
