## Supplemental Information for "HUH-tag conjugation enables accessible, site-specific functionalization of DNA origami with signaling-competent proteins"

**Contents:**

Table 1 - 9

**Table 1. Reagents and Tools.**

| Reagent or resource | Source | Identifier |
|---|---|---|
| **Experimental Models: Cell Lines** | | |
| Mouse: EL4.NOB-1 | ECACC | 87020408 |
| Mouse:EL4-IRAK1-mStaygold/mScarlet-NEMO | This paper | - |
| **sgRNA sequences** | | |
| IRAK1 (ENSMUSG00000031392) | 5’-TTAGAACCTGAAAAGAGCCA-3′ | - |
| Ikbkg (NEMO) (ENSMUSG00000004221) | 5′-GTTGGATGAACAAGCACCCC-3′ | - |
| **Recombinant DNA** | | |
| pMK-IRAK1-mStaygold-HDRtemp | This paper | - |
| pMK-_PuroR-2A-mScarlet-i3-NEMO HDR | This paper | - |
| pET28a-His10-mStayGold | This paper | |
| pET28a-His10-mScarlet-i3 | This paper | |
| pET28a-His10-IL-1β-HUH | This paper | |
| pMK | Thermo | GeneArt vector |
| pX330-U6-Chimeric_BB-CBh-hSpCas9 | Addgene | Plasmid #42230 |
| **Antibodies** | | |
| Rabbit monoclonal anti-IRAK1 (D51G7) | Cell Signaling Technology | Cat# 4504 |
| Rabbit polyclonal anti-IKK gamma/NEMO | abcam | Cat# ab244244 |

**Table 2. List of staple sequences for plate 1 of origami folding reaction.**

| Well | Sequence |
|---|---|

| | |
|---|---|
| A1 | CAGACGAAAAAGAAAGACTGGATAGCGTAGGCTTGAATACGTAATGCCACTACGTTT |
| A2 | GGTGGCACAATAAAAAGCAATACCAAAAAGCCTTTCTCATATATTTTAAATGCATTT |
| A3 | ATTTTCACATAGTTGTTCCGAAATCGAGCGGATTGCATCAAATTATAGTCAGAAGC |
| A4 | TACCGATTCGTCACCAGGAACGGTACTAATAGTAAAATGTTTGTTTTGCCAGAGGG |
| A5 | GAGGCGAAATATACACAATATAGAGATAGAACCCTGATAGCCCTAAAACACCTCAA |
| A6 | GCGAACTTCTGACCTGGTAATGCAATACACGAGCACTGCGCGTCACCCAGAACGTG |
| A7 | TACCGCCTCACGCATCCTCGTCTGGCAAGGGTCGAGAACAAGGCAGCAAAACGCGC |
| A8 | TCACCGTAGGGAAGATAAAGGGACTCCTTGTGTAGGTAAAGATAGAACCATTTCAA |
| A9 | CCGCCTGTGCGTATTCACAATCCCCGGGCGGTGCCACATCCCCACCGTCCATCCTC |
| A10 | AAGATTATTTAATTCTCCAACCTTTTGATAATTGCATATGCATATAACAGTTGATT |
| A11 | AGTCGGGTGAGCTAGGGGGTTTGGTGCTTATGAGCTCATTGCTTGCCGTCACAGGC |
| A12 | ATTTGCCTGAGAGAATGTGCTGCGCCATCGTGGGAGCCATCAACGGTAATCGTAAA |
| B1 | AGAGCCACAGGAGGCATTCCAACTAAAGTACGGTGTCCCGCCGGGCGCGGTTGCGG |
| B2 | TTTGAGCAAGAAACAATGATTAAGCCTGAGCGATGTTGGGAAGGGCGATCGGTTT |
| B3 | TTTCGTCAAAAATGAAAATACGATTTCGCTATTGGATAGCTCTCACGGAAAATTT |
| B4 | TTTGCCAAAAGGAATTACGAATGCAGAAGGGAATCAGTGAATAAGGCTTGCCTTT |
| B5 | TTTAGCGAGAGGCTTTTGCCGATAAATAAAACGTAGCCGGAACGAGGCGCAGTTT |
| B6 | TTTAAATCAGGTCTTTACCAATGACCTAATAATGCCCACGCATAACCGATATTTT |
| B7 | TTTACTTCAAATATCGCGTAGAGGAAAACTACAAATAGAAAGGAACAACTAATTT |
| B8 | TTTGTACCTTTAATTGCTCAGGTCAGGATATAATACCGTAACACTGAGTTTCTTT |
| B9 | TTTGCTCAACATGTTTTAATGAATATGGGGTCATACCAGGCGGATAAGTGCCTTT |
| B10 | TTTAAGCCTTAAATCAAGACTTGCGGACAGCGGGTAGAACGTCAGCGTGGTGTTT |
| B11 | TTTGGGCGCGAGCTGAAAAGCTATATTTCATCGCAGAGCCGCCACCAGAACCTTT |
| B12 | TTTAAGAATTAGCAAAATTTCATACATGAATTAGTTTGCCTTTAGCGTCAGATTT |
| C1 | TTTATACTTTTGCGGGAGAACATTATTACATACGTAAATATTGACGGAAATTTTT |
| C2 | TTTAAACCAAGTACCGCACTCCAAGAGCAGCAACCGCAAGCGGACTTATCAAAC |
| C3 | ACAAAGTCCCTGAAAGGTCACTCCGGCACCGCTTCACGCCAGGGTTTTC |
| C4 | TCTTACCAGATAACGATTCTCTCGCCATTCAGGCTCTGGCGAAAGGGGG |
| C5 | TTGAGAAATAATTAAACATACGGGGAGAGGCGGTTGCCCTGAGAGAGTT |
| C6 | TAAGGCGCTATATGACGCTGGGTTGTTCCAGTTTGGGTGCCGTAAAGCA |
| C7 | TGACCTAACGCGAGCCCTTCAGACTCCAACGTCAACACTACGTGAACCA |
| C8 | TTTTAACCCTTGAATTTTTTGGTGTAGCGGTCACGCGTATAACGTGCTT |
| C9 | ACATAAAACATTTATGCTTTGTTCTTTGATTAGTAACTATCGGCCTTGC |
| C10 | GAAGCGCCAAAATAGATTAAGAGTCCCGGAATTTGGCCAGCAGTTGGGC |
| C11 | ATTGTGTGATGAACGGTCAGTATTAAATTTAGGAATACCACAAGATTCA |

| | |
|---|---|
| C12 | TGCTCATCCGAACTTGTTACTAAAGAGGCGGGTAACAGGGAGAACCATC |
| D1 | ACAAAGCTAAATTGAAAAATCTACGTTAGGTAGAATTCAACTAGGCATA |
| D2 | GAAAAACCCGAGTAGAGCTAAAAAGGAGCTAAATCGTTGAGTTTTGCCC |
| D3 | AGCCATTGCAACAGAAAAGGGACATTCTTTAAAAATGATTATCAGATGA |
| D4 | GAGCGTCAATCAGAACATAAATTTCGTCTCGTCGCCAGCTTACGGCTGG |
| D5 | GCACCCAGCGTTTTTCTGCTCATAACGGAACGTGCAATGCCAACGGCAG |
| D6 | TCCGTTTAAAATCCCGGCGAACCAGTCACCAGCTTGTTGGTGTAGATGG |
| D7 | TGGCAGCGGTTGTGGTTTACCTTGGGTATGGTGCCGACCGTACATTTTT |
| D8 | GTAGGAACATGTAGCCATCCCTTTGCTCGTCATAAGGTGCCCCCTGCAT |
| D9 | AAGAAAAGTAATTTCAGTGTCTCTTCGCGTCCGTGAAGCATAAAGTGTA |
| D10 | TGCAGAAATAAAGTCAGCCAGTACCGAGCTCGAATAAATTGTTATCCGC |
| D11 | TTCAGCGCGTTGAAGTTCAGAGAATCCCCCTCAAATGAAAGCCGG |
| D12 | CATTAAACAAAAGACGTTTACGTAAGAGCAACACTATAATGGATT |
| E1 | ATAGTGGAGCCGCCACGGGAACGGGCCTTTCATCTTTTCATAAT |
| E2 | TGAAAGCGTAAGAATTAGTCTTTTGGATTATACTTCTGAATTT |
| E3 | TAACCACCACACCCCTATGGTACAATTTCATTTGAATTACTTT |
| E4 | TGGGCGCCAGGGTGCTGATTGAAAACTTTTTCAAATATATTTT |
| E5 | GAATACCCAAAGACGCCAGTTTGAGGAAATATTTAAATTGTA |
| E6 | CGAGGAATTATTTTGCGCATCAGATCGCACTCCAGCGACGTT |
| E7 | ATTAAGACACCCTCTAATGAGAAACCTGTCGTGCCCAGCAGG |
| E8 | ACCCTCAAAGTTTTCGAAAATTAGCCCGAGATAGGGGAACCC |
| E9 | TGAATTTATTGTATTAAAGGGAAGGGAAGAAAGCGACAGGAG |
| E10 | TTTTTCAGAGTGAGACGCCTGACCCATGGTATAGCTGCTCAG |
| E11 | TGAATTTGACAGCAGCCGATTAATCAGTGAGGCCAGCTCATG |
| E12 | CAGAGGCTATACCAGAAATACACCAGTCACACGACCCAGCAG |
| F1 | TGGTTTACAGTAGCGTAAAACTCACCGGAAACAATCGTAAAA |
| F2 | TTCATTATAATTTCACCAGTCAGGACGTAGCACCGCCTGCAA |
| F3 | CCTTAACATTTGAGGATTTAGGCCGTCAATAGATAATTGCGA |
| F4 | GTGTTGACGCTCAATCGTCTGACAGGGCCAGAATCCTGAGAA |
| F5 | TTTTTATAAAGGGAAGAAAGGAGCCCCCAAAAGAACCTGTTT |
| F6 | GATTTAGAGCTTGACGGGGCTAAGCAAAATCCCTTATAAATC |
| F7 | AGCTGCAAAGCCTGTGCCTGTACTGCGCCCTGCGGAGGTGTC |
| F8 | ACTCACATTAATTGCGTTGCCTGCCGTTTTCACGGTCATACC |
| F9 | GATAGCACGTTTGCAGTGATGAAGGGGCAAATGGTCAATAAC |
| F10 | AACGTCACAAAATCAAAGCCGTCCGGCAAACGCGGCAGCATC |
| F11 | AGGCGCTTTCGCACTCAATTGTCTAAAGTTAAACGATGCTGA |

| | |
|---|---|
| F12 | AGTGCCAAGCTTTCAGAGGTATAGGACGACGACAGTATCGGC |
| G1 | TTCAAAAGGGTGAGAAAGGCCGTATAAGCAAATAAAAATTTT |
| G2 | ACCGCCTAAACAAAAGCGGGGCGGGTCACTGTTGCGCCTGTG |
| G3 | ACCGTTCCAGTTAAGAATGCGGCGGGCGGATGGCTTAGAGCT |
| G4 | GAAAGCGTTCGGAACACTCTGTCTGCCAGCACGCGGGGTGCC |
| G5 | GTGCCTTTTTGATGGCATTGACCACCCTGCATTTTGAATCAA |
| G6 | GGGGTTTCCGGAATAAGCAAACGAGCTTCAAAGCGAACGCT |
| G7 | TTTCGGAATCGTCATAAATATTCATTAAACGAGCTGACTA |
| G8 | TTTTATTTTTGAATGGCTATACGTGGCACAGACAATTT |
| G9 | TTTGAGTAGAAGAACTCAAATAACATCACTTGCCTTTT |
| G10 | TTTCGCTACAGGGCGCGTAGCCGCGCTTAATGCGCTTT |
| G11 | TTTTATCAGGGCGATGGCCAGGGCGAAAAACCGTCTTT |
| G12 | TTTGTGAGACGGGCAACAGGTTTTTCTTTTCACCATTT |
| H1 | TTTAGCTGTTTCCTGTGTGTCGTAATCATGGTCATTTT |
| H2 | TTTGGCATCAGATGCCGGGTCAGCAAATCGTTAACTTT |
| H3 | TTTACGACGACAATAAACAAAGTAATTCTGTCCAGTTT |
| H4 | CACTGCCCGCTTTCCGATGGTGAGCGTAACGATCTA |
| H5 | AAGCAGAAAATTAATGCCGGAACTAGCATAACCAA |
| H6 | ACGCAATGTCAAATCACCATCAGCCCCAGTTAAAA |
| H7 | ATCGTCGAAAGAAGAGAGCGGAAAGAGTCTGTCCA |
| H8 | AAGAAACACAAACAACTAACAACTAATAGATTAGA |
| H9 | ACATTATATTAAATATCTAAAATATCTTACCCTCA |
| H10 | AATCTTGTGAATTATTTTAAGAACTGGCTCATTAT |
| H11 | AATTAACCGTTGTAATCCAGAAGTAACAGTACCTT |
| H12 | CGGGCGCTAGGGCGTAGAATCATGATGAAACAAAC |

**Table 3. List of staple sequences for plate 2 of origami folding reaction.**

| Well | Sequence |
|---|---|
| A1 | AGTCCACTATTAAAAATCAAGAACATAGCGATAGC |
| A2 | TTAATGAATCGGCCGCGGTCCTAAATGCTGATGCA |
| A3 | GAGCCGGAGCCTCCCAGACGAAGGTTTCACGCAAC |
| A4 | TCACAGTTGAGGATTCCACACCTAGAAAAAGCCTG |
| A5 | TAAGAGGTCATTTTAGACCGGAGGTGTATCACCGT |
| A6 | CTGGTAATGGGTAATCCAGCGAGGCAGAGGCATTT |
| A7 | TTACACTGGTGTGTTTACCTGACCGACAAAAGGTA |

| | |
|---|---|
| A8 | CTCCGGCCAGAGCAGGTGGTGAAACCAATCAATAA |
| A9 | CCATTAGATACATTGAAGTTTTTGAGGCAGGTCAG |
| A10 | ACGTACAGCGCCATTACATCGTATAGAAGGCTTAT |
| A11 | TAGACTTTCTCCGTTTAAATTAGCGAACCTCCCGA |
| A12 | GGTGAAGACGCCAGGCGCAACGTAACAACTGGCCT |
| B1 | GATAACCGACGGCCCTCAGGAGTAACCGATATTTT |
| B2 | GAGGGTAGCTATTTTTGAGAGTCGATGAAAAATAA |
| B3 | AATATGATATTCAACCGTTCTACCCCGGTTGTTAA |
| B4 | TTGAGGGCACCGACTAACATCTCAATTCTACTA |
| B5 | TTTGCGAACGAGTAGATTTAGTTTGACTGTTTA |
| B6 | ATTTACATTGGGTGAGGCGGTGTACAGACCAG |
| B7 | CGAACGTGGCGTTTTAGACCTCAGCAGCGAAA |
| B8 | TTTTTTAGTTAATTTCGTTATACAAATTTT |
| B9 | TTTCTTTTTTAATGGTGAGAAGAGTCATTT |
| B10 | TTTTAATGGAAGGGTACAATAACGGATTTT |
| B11 | AATAGCAAAGGCTATCAGGTCATTGCTTT |
| B12 | GCCGCCAATACAGGAGTGTACTGGTATTT |
| C1 | ATTGCGTATATTCCTACCGAATCTAAAG |
| C2 | TACCATACTGATTGTTAATGCATCAATA |
| C3 | ATTTGTAGCGCATAAAGATAAGAGCCAG |
| C4 | AGGCAAAGCAAGGCAACAGCCATATTAT |
| C5 | TTTAAACGTAGAAAAGACCCTGTATTT |
| C6 | TTTGTCGAGAGGGTTGATTAGAGATTT |
| C7 | TTTGTCACCAGTACAGCCCGAAAGTTT |
| C8 | TTTAGGAATTGCGAAATAAATCAATTT |
| C9 | TTTATTCGGTCGCTGCCAATACTGTTT |
| C10 | TTTAAGGCACCAACCAACCAAAATTTT |
| C11 | TTTACGGTCAATCATATACATAACTTT |
| C12 | TTTCTGACGAGAAACGAACTAACGTTT |
| D1 | TTTATTCATTAAAGGGGCAAGGCATTT |
| D2 | TTTCTGGTCTGGTCAACGGGTATTTTT |
| D3 | TTTAGAGACGCAGAAGAGGTTTTGTTT |
| D4 | TTTTGCGGGCCTCTTTTTGTTTAATTT |
| D5 | TTTCAACATTAAATGCAATAATAATTT |
| D6 | TTTCTGTAGCGCGTTTTTCATTTGTTT |
| D7 | TTTACCACCAGAGCCCCCAATTCTTTT |

| | |
|---|---|
| D8 | TTTATAAGTTTTAACAATGCTGTATTT |
| D9 | TAACCCTATACACTAAAACAC |
| D10 | TTAAACAAATCTCCAAAAAAA |
| D11 | GCGGCCATGCCCCCTGCCTAT |
| D12 | GTAGCATTTGAGCCATTTGGG |
| E1 | /5Biosg/TTTTCTGGTCGAAGGTTCCTTTGCCCGAACGTTATTTTT |
| E2 | /5Biosg/TTTCAGTGCCACGCTGAAACAGAGCAGATTCCTACATT |
| E3 | /5Biosg/TTTCGCAAGGGCTAAATCGGTTGTAAAGCCTCAGAGCA |
| E4 | /5Biosg/TTTCAGCAAATGAAAAACGAACCACAGTAAT |
| E5 | /5Biosg/TTTCATCACCTTGCTGAATCGCCAGGCCAAC |
| E6 | /5Biosg/TTTATATCAATAGGAGCATTCGACAACTCGT |
| E7 | /5Biosg/TTTTCAGTTGTGGGAAGGGCTTGAGATGGTT |
| E8 | /5Biosg/TTTTTCGCATTAAATTTTTGATAATCAGAAA |
| E9 | /5Biosg/TTTATCAGCTATGGGATCAAAGTCAGAGGGT |
| E10 | /5Biosg/TTTTAGGAACACAAACGGCGGATTGGAAACC |
| E11 | /5Biosg/TTTTTCGCGTCCCGTCGCCACAAGAATTGAG |
| E12 | /5Biosg/TTTAACGTTATGCATCTACCACGGAATAAGT |
| F1 | TTTGAACAACATTATTACAATAAAACACCAGAACGAGTAG |
| F2 | TTTGTTGAAAGGAATTGAGAGTTGGCAAATCAACATTT |
| F3 | TTTCTGAGAGTCTGGTCCTGTAGCCAGCTTTCATTTT |
| F4 | TTTATGCCTGAGTAATATTACGCAGTATGTTAGCTTT |

**Table 4. List of handle-free staple sequences for plate 3 of origami folding reaction.**

| Well | Sequence |
|---|---|
| A1 | CGACATTAGAAACGCAAAAGAACTGGCA |
| A2 | AAAACAGGAAGATTGGAGACAAATAACG |
| A3 | GTCACAATCAATCATACCAGAAGGAAAC |
| A4 | TGTCAATCATATGTAGCTGATTAGCCGA |
| A5 | AACATAAATCAGAGGAAGCCCTTTTTAA |
| A6 | AGCAAACAAGAGAAATCTACAATAGCTA |
| A7 | TGATTAATGGCAACATATAAACAACCGA |
| A8 | CCAATGAAAATCACCCAGCGCCAAAGAC |
| A9 | TTAACTGAAAGAAAATTCATA |
| A10 | TTACCAACCAGTTAATTAGACGGGAGAA |
| A11 | GAAAAGTAATTGAGCGCTAATAAACAGG |

| | |
|---|---|
| A12 | TTAGTTGATAAGAAAGCAGCCTTTACAG |
| B1 | GAACCGCTTATTAGGCACCGTAATCAGT |
| B2 | AAAAGGGAATTAGAGCCAGCAAACCATC |
| B3 | ACCGGAACCAGACATTAGCAAGGCCGGA |
| B4 | ACCATTACCATTTCCAGAGCCTAATTTGCGCTAAC |
| B5 | TTTTTATACGCGAGGCTACAATTTTATC |
| B6 | AGAGAATTTATCCCAATCCAACTATTTT |
| B7 | AGCGACACGGTCATAGCCCCCCACCCTC |
| B8 | CAGTCTCTATTCACCCCTCAGAGCCGCC |
| B9 | AATAGCAAGGCCACCACCGGA |
| B10 | GATAAGTTTACGAGTCATTACCGCGCCC |
| B11 | CTGAATCCCGGTATTCTAAGATTTCATC |
| B12 | ACATGTTTTATCATTCATCGAGAACAAG |
| C1 | GGATTAGGTATAAACAGTAAGCGTCATA |
| C2 | ACCCTCAACGATTGGCCTTGATGAATTT |
| C3 | CCTATTATTCTGATATAAAGCCAGAATG |
| C4 | TAAATCCTCATTAATATCCCATCCTAATCCTGAAC |
| C5 | ACAGTAGAGAGAATCGCGCCTGTTTATC |
| C6 | CAAGCCGTCGGCTGTCTTTCCCAGCTAA |
| C7 | CATGGCTGAGTAACAGTGCCCGATTAGC |
| C8 | GAGCCACGTACCGCGGCTGAGACTCCTC |
| C9 | AACGCCAACAAACATGAAAGT |
| C10 | GACCGTGCGGAATCTCGCCATATTTAAC |
| C11 | AACAATATCGAGCCAGTAATAGGCTTAA |
| C12 | TTTCTTACCAGTATAAAGCCA |
| D1 | CAACTTTCAGCCCTGGGATAGCAAGCCC |
| D2 | AAGAGAAACTCAGGAGGTTTACACCCTC |
| D3 | GTCGTCTTTCCAAATTCTCAGAACCGCC |
| D4 | AGAACCGCCACCAAATAAGAATAAACACTGATAAA |
| D5 | CTGAGAGACAAAGAAATTTAATGGTTTG |
| D6 | ACGCTCATTTAGTATCATATGCATCTTC |
| D7 | AATAGGATAGCATTCCACAGACAACAGT |
| D8 | CTTAAACGCCTTTATCTGTATGGGATTT |
| D9 | GGGTTATATGACGTTAGTAAA |
| D10 | CCTTGCTTTAGAATCTCCGGCTTAGGTT |
| D11 | AAATACCAATCCAATCGCAAGACTACCT |

| | |
|---|---|
| D12 | TTTATAGTGAATTTATCAAAAT |
| E1 | CATGAGGTGCGGGAAGTTGCGCCGACAA |
| E2 | TGCTAAAAGGCTCCAAAAGGAAGCTTGA |
| E3 | TCGGAACGAGGGCACTTTGCTTTCGAGG |
| E4 | CGGTTTATCAGCATTAATTAATTTTCCCTCTGTAA |
| E5 | TACAAAAATTAATTTCAATATATGTGAG |
| E6 | CATAGGTTTAGATTAAGACGCAAACAGT |
| E7 | TGACAACTTAAAGGCCGCTTTAAGTTTC |
| E8 | TCATCGCCAGCGATTTTGAGGACTAAAG |
| E9 | TTACCTGAGTAGCAACGGCTA |
| E10 | ACAGAAATCAGATGATTATTCATTTCAA |
| E11 | TGAATAAATCAAGAAAACAAATCGCGCA |
| E12 | TTTTCGCCTGATTGCTTTGAAT |
| F1 | CCCAAATGAGGACACGAAATCCGCGACC |
| F2 | ACTTTTTTCATCTTTGACCCCCTGATAA |
| F3 | GGCTGGCTGACCTCAGAGTACAACGGAG |
| F4 | AGCGCGAAACAAATTTTCAGGTTTAACGTAAAGAA |
| F5 | CATTTTGTATAATCTCAAAATTATTTGC |
| F6 | ACCAAGTTTACATCGGGAGAATAGAACC |
| F7 | TGCTCCAGACCAACTTTGAAACAACGTA |
| F8 | AACTTTAATCATTGACAAGAACCGGATA |
| F9 | GAATTATCATTCATCAAGAGT |
| F10 | AAGTATTAGACTTTCACCAGAAGGAGCG |
| F11 | ACGTAAATGGCAATTCATCAACGGAACA |
| F12 | TTTAATTTTAAAAGTTTGAGTA |

**Table 5. List of staple sequences containing attachment sequence (highlighted yellow) L1 for plate 3 of origami folding reaction.**

| Well | Sequence |
|---|---|
| A1 | CGACATTAGAAACGCAAAAGAACTGGCATTTTTTTT**CCCTCATTCAATACCCTAGG** |
| A2 | AAAACAGGAAGATTGGAGACAAATAACGTTTTTTTT**CCCTCATTCAATACCCTAGG** |
| A3 | GTCACAATCAATCATACCAGAAGGAAACTTTTTTTT**CCCTCATTCAATACCCTAGG** |
| A4 | TGTCAATCATATGTAGCTGATTAGCCGATTTTTTTT**CCCTCATTCAATACCCTAGG** |
| A5 | AACATAAATCAGAGGAAGCCCTTTTTAATTTTTTTT**CCCTCATTCAATACCCTAGG** |

| | |
|---|---|
| A6 | AGCAAACAAGAGAAATCTACAATAGCTATTTTTTTT**CCCTCATTCAATACCCTAGG** |
| A7 | TGATTAATGGCAACATATAAACAACCGATTTTTTTT**CCCTCATTCAATACCCTAGG** |
| A8 | CCAATGAAAATCACCCAGCGCCAAAGACTTTTTTTT**CCCTCATTCAATACCCTAGG** |
| A9 | TTAACTGAAAGAAAATTCATATTTTTTTTT**CCCTCATTCAATACCCTAGG** |
| A10 | TTACCAACCAGTTAATTAGACGGGAGAATTTTTTTT**CCCTCATTCAATACCCTAGG** |
| A11 | GAAAAGTAATTGAGCGCTAATAAACAGGTTTTTTTT**CCCTCATTCAATACCCTAGG** |
| A12 | TTAGTTGATAAGAAAGCAGCCTTTACAGTTTTTTTT**CCCTCATTCAATACCCTAGG** |
| B1 | GAACCGCTTATTAGGCACCGTAATCAGTTTTTTTTT**CCCTCATTCAATACCCTAGG** |
| B2 | AAAAGGGAATTAGAGCCAGCAAACCATCTTTTTTTT**CCCTCATTCAATACCCTAGG** |
| B3 | ACCGGAACCAGACATTAGCAAGGCCGGATTTTTTTT**CCCTCATTCAATACCCTAGG** |
| B4 | ACCATTACCATTTCCAGAGCCTAATTTGCGCTAACTTTTTTTT**CCCTCATTCAATACCCTAGG** |
| B5 | TTTTTATACGCGAGGCTACAATTTTATCTTTTTTTT**CCCTCATTCAATACCCTAGG** |
| B6 | AGAGAATTTATCCCAATCCAACTATTTTTTTTTTTTT**CCCTCATTCAATACCCTAGG** |
| B7 | AGCGACACGGTCATAGCCCCCACCCTCTTTTTTTT**CCCTCATTCAATACCCTAGG** |
| B8 | CAGTCTCTATTCACCCCTCAGAGCCGCCTTTTTTTT**CCCTCATTCAATACCCTAGG** |
| B9 | AATAGCAAGGCCACCACCGGATTTTTTTT**CCCTCATTCAATACCCTAGG** |
| B10 | GATAAGTTTACGAGTCATTACCGCGCCCTTTTTTTT**CCCTCATTCAATACCCTAGG** |
| B11 | CTGAATCCCGGTATTCTAAGATTTCATCTTTTTTTT**CCCTCATTCAATACCCTAGG** |
| B12 | ACATGTTTTATCATTCATCGAGAACAAGTTTTTTTT**CCCTCATTCAATACCCTAGG** |
| C1 | GGATTAGGTATAAACAGTAAGCGTCATATTTTTTTT**CCCTCATTCAATACCCTAGG** |
| C2 | ACCCTCAACGATTGGCCTTGATGAATTTTTTTTTTT**CCCTCATTCAATACCCTAGG** |
| C3 | CCTATTATTCTGATATAAAGCCAGAATGTTTTTTTT**CCCTCATTCAATACCCTAGG** |
| C4 | TAAATCCTCATTAATATCCCATCCTAATCCTGAACTTTTTTTT**CCCTCATTCAATACCCTAGG** |
| C5 | ACAGTAGAGAGAATCGCGCCTGTTTATCTTTTTTTT**CCCTCATTCAATACCCTAGG** |
| C6 | CAAGCCGTCGGCTGTCTTTCCCAGCTAATTTTTTTT**CCCTCATTCAATACCCTAGG** |
| C7 | CATGGCTGAGTAACAGTGCCCGATTAGCTTTTTTTT**CCCTCATTCAATACCCTAGG** |
| C8 | GAGCCACGTACCGCGGCTGAGACTCCTCTTTTTTTT**CCCTCATTCAATACCCTAGG** |
| C9 | AACGCCAACAAACATGAAAGTTTTTTTTT**CCCTCATTCAATACCCTAGG** |
| C10 | GACCGTGCGGAATCTCGCCATATTTAACTTTTTTTT**CCCTCATTCAATACCCTAGG** |
| C11 | AACAATATCGAGCCAGTAATAGGCTTAATTTTTTTT**CCCTCATTCAATACCCTAGG** |
| C12 | TTTTCTTACCAGTATAAAGCCATTTTTTTTT**CCCTCATTCAATACCCTAGG** |
| D1 | CAACTTTCAGCCCTGGGATAGCAAGCCCTTTTTTTT**CCCTCATTCAATACCCTAGG** |
| D2 | AAGAGAAACTCAGGAGGTTTACACCCTCTTTTTTTT**CCCTCATTCAATACCCTAGG** |
| D3 | GTCGTCTTTCCAAATTCTCAGAACCGCCTTTTTTTT**CCCTCATTCAATACCCTAGG** |
| D4 | AGAACCGCCACCAAATAAGAATAAACACTGATAAATTTTTTTT**CCCTCATTCAATACCCTAG** **G** |

| | |
|---|---|
| D5 | CTGAGAGACAAAGAAATTTAATGGTTTGTTTTTTTT**CCCTCATTCAATACCCTAGG** |
| D6 | ACGCTCATTTAGTATCATATGCATCTTCTTTTTTTT**CCCTCATTCAATACCCTAGG** |
| D7 | AATAGGATAGCATTCCACAGACAACAGTTTTTTTT**CCCTCATTCAATACCCTAGG** |
| D8 | CTTAAACGCCTTTATCTGTATGGGATTTTTTTTTT**CCCTCATTCAATACCCTAGG** |
| D9 | GGGTTATATGACGTTAGTAAATTTTTTTTT**CCCTCATTCAATACCCTAGG** |
| D10 | CCTTGCTTTAGAATCTCCGGCTTAGGTTTTTTTTT**CCCTCATTCAATACCCTAGG** |
| D11 | AAATACCAATCCAATCGCAAGACTACCTTTTTTTTT**CCCTCATTCAATACCCTAGG** |
| D12 | TTTATAGTGAATTTATCAAAATTTTTTTTT**CCCTCATTCAATACCCTAGG** |
| E1 | CATGAGGTGCGGGAAGTTGCGCCGACAATTTTTTTT**CCCTCATTCAATACCCTAGG** |
| E2 | TGCTAAAAGGCTCCAAAAGGAAGCTTGATTTTTTTT**CCCTCATTCAATACCCTAGG** |
| E3 | TCGGAACGAGGGCACTTTGCTTTCGAGGTTTTTTTT**CCCTCATTCAATACCCTAGG** |
| E4 | CGGTTTATCAGCATTAATTAATTTCCCTCTGTAATTTTTTTT**CCCTCATTCAATACCCTAGG** |
| E5 | TACAAAAATTAATTTCAATATATGTGAGTTTTTTTT**CCCTCATTCAATACCCTAGG** |
| E6 | CATAGGTTTAGATTAAGACGCAAACAGTTTTTTTT**CCCTCATTCAATACCCTAGG** |
| E7 | TGACAACTTAAAGGCCGCTTTAAGTTTCTTTTTTTT**CCCTCATTCAATACCCTAGG** |
| E8 | TCATCGCCAGCGATTTTGAGGACTAAAGTTTTTTTT**CCCTCATTCAATACCCTAGG** |
| E9 | TTACCTGAGTAGCAACGGCTATTTTTTTT**CCCTCATTCAATACCCTAGG** |
| E10 | ACAGAAATCAGATGATTATTCATTTCAATTTTTTTT**CCCTCATTCAATACCCTAGG** |
| E11 | TGAATAAATCAAGAAAACAAATCGCGCATTTTTTTT**CCCTCATTCAATACCCTAGG** |
| E12 | TTTTCGCCTGATTGCTTTGAATTTTTTTTT**CCCTCATTCAATACCCTAGG** |
| F1 | CCCAAATGAGGACACGAAATCCGCGACCTTTTTTTT**CCCTCATTCAATACCCTAGG** |
| F2 | ACTTTTTTCATCTTTGACCCCCTGATAATTTTTTTT**CCCTCATTCAATACCCTAGG** |
| F3 | GGCTGGCTGACCTCAGAGTACAACGGAGTTTTTTTT**CCCTCATTCAATACCCTAGG** |
| F4 | AGCGCGAAACAAATTTTCAGGTTTAACGTAAAGAATTTTTTTT**CCCTCATTCAATACCCTAG**<br>**G** |
| F5 | CATTTTGTATAATCTCAAAATTATTTGCTTTTTTTT**CCCTCATTCAATACCCTAGG** |
| F6 | ACCAAGTTTACATCGGGAGAATAGAACCTTTTTTTT**CCCTCATTCAATACCCTAGG** |
| F7 | TGCTCCAGACCAACTTTGAAACAACGTATTTTTTTT**CCCTCATTCAATACCCTAGG** |
| F8 | AACTTTAATCATTGACAAGAACCGGATATTTTTTTT**CCCTCATTCAATACCCTAGG** |
| F9 | GAATTATCATTCATCAAGAGTTTTTTTTT**CCCTCATTCAATACCCTAGG** |
| F10 | AAGTATTAGACTTTCACCAGAAGGAGCGTTTTTTTT**CCCTCATTCAATACCCTAGG** |
| F11 | ACGTAAATGGCAATTCATCAACGGAACATTTTTTTT**CCCTCATTCAATACCCTAGG** |
| F12 | TTTAATTTTAAAAGTTTGAGTATTTTTTTT**CCCTCATTCAATACCCTAGG** |

**Table 6. List with ATTO647N-containing staples for flexible fluorescent labeling of origami structures**

| Staple name | Sequence |
|---|---|
| 157+dye | /5ATTO647NN/TTTCTGAGAGTCTGGTCCTGTAGCCAGCTTTCATTTT |
| 158+dye | /5ATTO647NN/TTTATGCCTGAGTAATATTACGCAGTATGTTAGCTTT |
| 159+dye | /5ATTO647NN/TTTGTTGAAAGGAATTGAGAGTTGGCAAATCAACATTT |
| 160+dye | /5ATTO647NN/TTTGAACAACATTATTACAATAAAACACCAGAACGAGTAG |

**Table 7. gBlocks used for cloning.**

| **Function** | **Sequence** | **Supplier** |
|---|---|---|
| gBlock for cloning of pET28a-His$_{10}$-IL1β-FBNYV into pET28a-IL1β-spytag | ACTTTAAGAAGGAGATATACCATGGGCAGCAGCCATCATCA<br>TCATCATCACCATCACCATCACGGTGGTGGTAGCGGTGGAG<br>GAGGATGGATGAGCGAGAATCTTTATTTTCAGGGCGGGGGT<br>GGCATGGTCCCGATTCGCCAACTTCACTATCGCCTTCGTGAT<br>GAGCAACAAAAATCGCTGGTCCTTTCTGATCCCTATGAGCTT<br>AAAGCCTTACACTTGAACGGTCAAAATATCAACCAGCAGGT<br>CATTTTTAGCATGTCCTTTGTGCAGGGTGAACCTTCTAACGA<br>TAAGATTCCGGTAGCGCTGGGGTTAAAAGGAAAAAATCTTT<br>ACCTGTCGTGTGTTATGAAGGATGGTACGCCGACTTTACAA<br>CTGGAGTCGGTTGACCCCAAGCAGTACCCTAAGAAGAAAAT<br>GGAGAAACGCTTCGTTTTCAACAAGATTGAAGTGAAGAGTA<br>AAGTCGAGTTTGAATCCGCCGAGTTCCCCAACTGGTACATTT<br>CTACTTCTCAAGCCGAGCATAAACCCGTATTTCTGGGGAATA<br>ACTCTGGGCAGGATATTATCGATTTCACGATGGAGAGTGTGT<br>CGTCAGGTGGAGGATCTGGGGGTGGCGGTTCCGGTGGTATG<br>GCGCGCCAGGTGATTTGCTGGTGCTTTACCCTGAACAACCC<br>GCTGAGCCCGCTGAGCCTGCATGATAGCATGAAATATCTGG<br>TGTATCAGACCGAACAGGGCGAAGCGGGCAACATTCATTTT<br>CAGGGCTATATTGAAATGAAAAAACGCACCAGCCTGGCGG<br>GCATGAAAAAACTGATTCCGGGCGCGCATTTTGAAAAACGC<br>CGCGGCACCCAGGGCGAAGCGCGCGCGTATAGCATGAAAG<br>AAGATACCCGCCTGGAAGGCCCGTGGGAATATGGCGAATT<br>TGTGCCGTAGACTAGTTAGCGGCCGCACTCGAGCA | **IDT DNA Technologies** |

**Table 8. List of oligonucleotides.**

| Function | Sequence | Supplier |
|---|---|---|

| | | |
|---|---|---|
| Attachment oligonucleotide for conjugation to HUH-tag | AAGTATTACCAGAAACCTAGGGTATTGAATGAGGG | IDT DNA Technologies |
| Attachment oligonucleotide for conjugation to HUH-tag and fluorescent labeling of target protein | AAGTATTACCAGAAACCTAGGGTATTGAATGAGGGTTT-Cy5 | IDT DNA Technologies |
| Attachment oligonucleotide for conjugation to HUH-tag and fluorescent labeling of target protein | AAGTATTACCAGAAACCTAGGGTATTGAATGAGGGTTT - Atto647N | IDT DNA Technologies |
| T7 promotor forward primer | TAATACGACTCACTATAGGG | IDT DNA Technologies |
| T7 terminator reverse primer | GCTAGTTATTGCTCAGCGG | IDT DNA Technologies |

**Table 9. Cost comparison of ssDNA reagents for HUH-tag versus conventional chemical tag-based protein-DNA conjugation**

*Prices reflect standard list prices for custom oligonucleotide synthesis at IDT, retrieved on 23/09/2026. Prices are per single oligonucleotide at a synthesis scale sufficient for typical labeling reactions (see Methods) and are given in Euro (€). Prices vary with oligonucleotide length, purification method, and scale; the values below use a representative 35-nt oligonucleotide for direct comparison.*

| Component | Vendor | Specification | Typical yield/size | Cost per order | Subtotal per nmol |
|---|---|---|---|---|---|
| **HUH-tag conjugation (this study)** | | | | | |
| ssDNA, unmodified | IDT | 35 nt, HPLC-purified, 100 nmol | 3.5 nmol (guaranteed) | € 37.10 (~$ 42.30) | € 10.60 (~$ 12.10) |
| **SNAP-tag / benzylguanine-modified ssDNA (self-made)** | | | | | |
| 5'-amino-modified ssDNA | IDT | 35 nt, 5' amino-C6, HPLC-purified, 100 nmol | 3.5 nmol (guaranteed) | € 57.90 (~$ 66.10) | € 16.50 (~$ 18.80) |
| BG-GLA-NHS coupling reagent | NEB | Catalogue number: S9151S | 2 mg | € 505.00 (~$ 576.60) | € 0.12 (~$ 0.14) |

| | | | | | |
|---|---|---|---|---|---|
| Subtotal per nmol conjugate (at 50X excess of BG-GLA-NHS) | | | | | € 22.50 (~$ 25.70) |
| **SNAP-tag / benzylguanine-modified ssDNA (synthesized)** | | | | | |
| 5'-BG-modified ssDNA | biomers | 35 nt, 5’-BG-modified, HPLC-purified, 0.05 µmol | 4.7 nmol (guaranteed only for a 20mer) | € 303.25 (~$ 346.20) | € 64.50 (~$ 73.70) |
